# Alanine Scanning to Define Membrane Protein-Lipid Interaction Sites Using Native Mass Spectrometry

**DOI:** 10.1101/2024.10.24.620105

**Authors:** Hiruni S. Jayasekera, Farhana Afrin Mohona, Madison J. De Jesus, Katherine M. Miller, Michael T. Marty

## Abstract

Lipids surrounding membrane proteins interact with different sites on the protein at varying specificities, ranging from highly specific to weak interactions. These interactions can modulate the structure, function, and stability of membrane proteins. Thus, to better understand membrane protein structure and function, it is important to identify the locations of lipid binding and the relative specificities of lipid binding at these sites. In our previous native mass spectrometry (MS) study, we developed a single and double mutant analysis approach to profile the contribution of specific residues toward lipid binding. Here, we extend this method by screening a broad range of mutants of AqpZ to identify specific lipid binding sites and by measuring binding of different lipid types to measure the selectivity of different lipids at selected binding sites. We complemented these native MS studies with molecular dynamics (MD) simulations to visualize lipid interactions at selected sites. We discovered that AqpZ is selective towards cardiolipins (CL) but only at specific sites. Specifically, CL orients with its headgroup facing the cytoplasmic side, and its acyl chains interact with a hydrophobic pocket located at the monomeric interface within the lipid bilayer. Overall, this integrative approach provides unique insights into lipid binding sites and the selectivity of various lipids towards AqpZ, enabling us to map the AqpZ protein structure based on the lipid affinity.

## Introduction

Membrane proteins are important drug targets and play major physiological roles, such as signaling, transport, and catalysis.^1–3^ Lipids in the surrounding bilayer can modulate the structure, function, and stability of membrane proteins.^4^ These membrane protein-lipid interactions range from highly specific interactions at particular binding sites to nonspecific, transient interactions.^5–7^ However, it is challenging to map the specificity of different lipids to different potential binding sites on membrane proteins.

Aquaporin Z (AqpZ, UniProt: P60844) is a water channel in *E. coli*.^8^ Previous studies have shown that AqpZ interacts with phosphatidylglycerol (PG) and cardiolipin (CL) lipids,^9–11^ and a study indicates that CL facilitates water transport activity.^12^ Molecular dynamic (MD) simulations have localized PG and CL interactions towards exposed cationic residues, primarily on the cytoplasmic surface.^11,13,14^ Specifically, Schmidt *et al*. observed PG is more uniformly distributed around the protein, but CL tends to associate more closely with interfaces between monomers.^11^ Corey *et al*. found that CL headgroups interacted more with Arg than Lys residues.^13^ In contrast, a recent cryo-EM study modeled CL to lipid-like density on the periplasmic side of the membrane.^15^ Thus, additional experimental evidence is needed to determine the specific binding sites and the site selectivity towards different lipids.

Native mass spectrometry (MS) has emerged as a valuable tool in studying membrane protein-lipid interactions. Native MS uses nondenaturing ionization conditions to preserve native protein conformations and noncovalent interactions for mass analysis.^16–18^ Previously, we developed a native MS approach to measure the thermodynamic contributions of specific amino acid residues on AqpZ toward CL binding by simultaneously analyzing a mutant protein and the wild type.^6^ We discovered that W14 residue contributes to the highest affinity binding site, and the R224 residue contributes to the second highest affinity binding site.^6^

Here, we extended these studies by measuring thermodynamic contributions of an expanded set of amino acid mutations (**Figure 1A–B**) to map CL binding hotspots. We then evaluated the selectivity of these CL binding sites for other lipids by comparing binding of four different lipid types (**Figure 1C–F**) with the mutants. To complement the native MS studies, we conducted a series of coarse-grain MD simulations of wild-type AqpZ in a model bacterial membrane.^19,20^ The occupancy and residence time analyses of different lipids at each residue were compared to characterize the selectivity of different potential lipid binding sites observed using native MS. Combining the native MS and MD results, we construct a map of lipid binding and selectivity to AqpZ.

**Figure 1.**
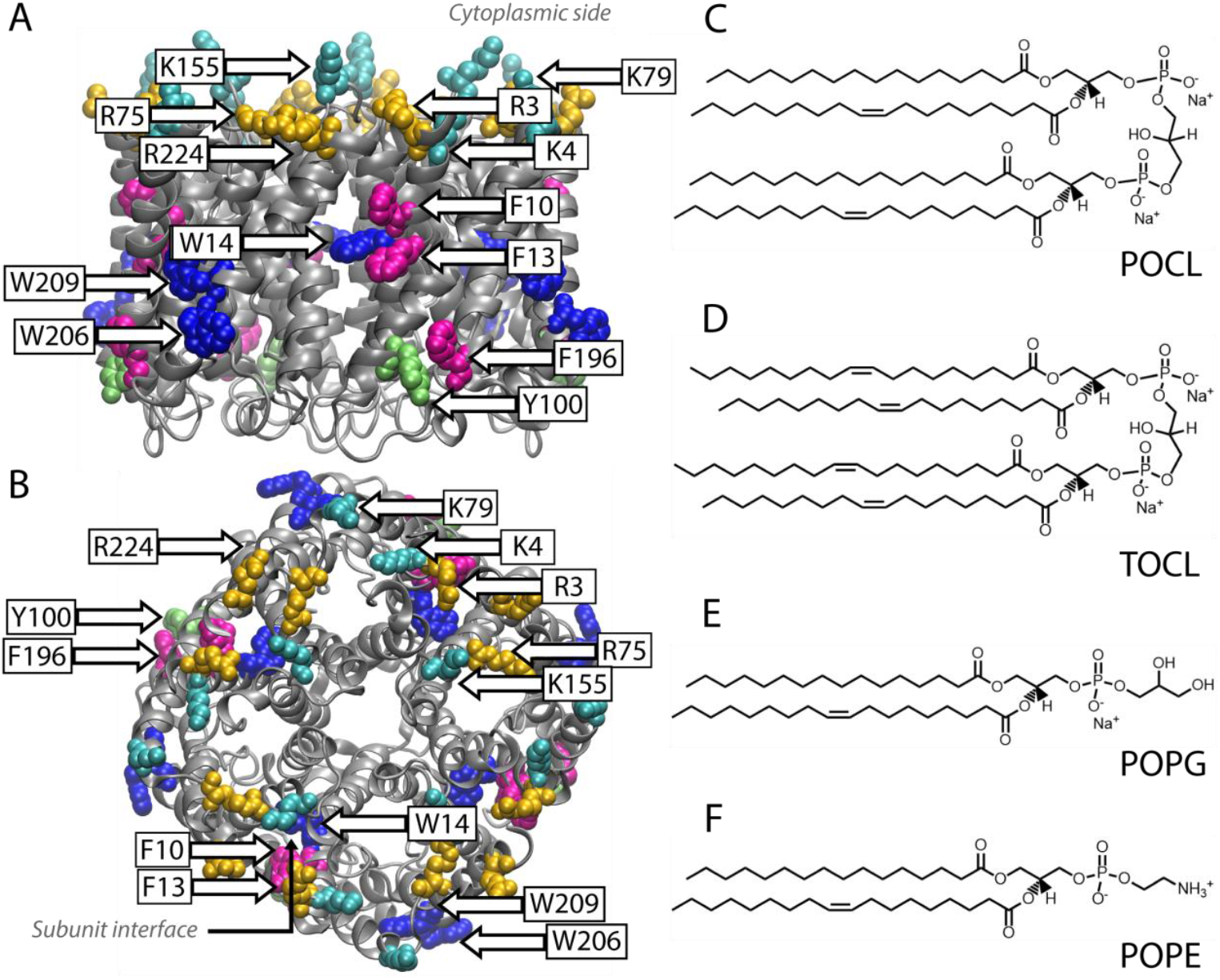
(A) Side-view of AqpZ with the cytoplasmic side of the membrane on top and (B) top-view of AqpZ (cytoplasmic face in front) with mutant sites W14, W206, and W209 (in *blue*); F10, F13, and F196 (in *magenta*); Y100 (*green*); R3, R75, and R224 (in *yellow*); and K4, K79, and K155 (in *cyan*) labeled, indicated with an arrow, and colored based on residue name. Four chains of the protein are shown in *gray* and one of the subunit interfaces is labelled with an arrow. The PDB code is 1RC2. Structures of four lipid types: (C) POCL, (D) TOCL, (E) POPG, and (F) POPE.

## Experimental Section

### AqpZ Mutagenesis, Expression, and Purification

We selected residues to mutate based on three criteria that included: 1) mutants from prior studies^6,11^ that are known to affect lipid binding (W14), 2) hydrophobic residues that likely interact with the lipid tails (F10, F13, F196, W206, W209, and Y100), and 3) cationic residues at the lipid interface that could interact with anionic CL or PG headgroups (R3, R75, R224, K4, K79, and K155), as shown in **Figure 1A, 1B**. After selecting these 13 residues, we designed primers for the alanine mutants and performed site-directed mutagenesis, as previously described.^6^ Alanine was selected as a neutral replacement residue that contrasted with the bulky hydrophobic residues and cationic residues.

The mutant plasmids were sequenced before transformation into *E. coli* OverExpress C43 (DE3) competent cells (Sigma Aldrich). We followed the previously described protocol for expressing and purifying AqpZ proteins.^6,12,21,22^ Following purification, all the proteins were exchanged into buffer containing 0.2 M ammonium acetate with 0.5% tetraethylene glycol monooctyl ether (C8E4).^6,11^ Purified and buffer exchanged individual proteins were analyzed using native MS, and we observed exclusively tetramers with similar charge state distributions to the wild-type protein, which indicates that the mutations did not significantly affect the AqpZ structure (**Figure S1**).

### Native MS Sample Preparation and Analysis

We tested binding of four lipids: 1′,3′-bis[1-palmitoyl-2-oleoyl-*sn*-glycero-3-phospho]-glycerol (POCL), 1’,3’-bis[1,2-dioleoyl-*sn*-glycero-3-phospho]-glycerol (TOCL), 1-palmitoyl-2-oleoyl-*sn*-glycero-3-phospho-(1’-rac-glycerol) (POPG), and 1-palmitoyl-2-oleoyl-*sn*-glycero-3-phosphoethanolamine (POPE). Their structures are shown in **Figure 1C–F**. Lipids were purchased from Avanti Polar Lipids, and lipid concentrations in chloroform were quantified using phosphate analysis.^6,11,12^ After drying off the chloroform, we prepared stock lipid solutions in 0.2 M ammonium acetate with 0.5% C8E4, and we performed another phosphate analysis for the accurate quantitation of the lipids in the detergent solutions.^6^ The stock concentrations were around 1–1.5 mM.

Single mutant analysis was performed as previously described.^6^ Briefly, we mixed pairwise combinations of wild-type protein with each mutant approximately at a 1:1 molar ratio. We slightly adjusted the protein ratio to achieve roughly equal intensities of the tetramer on the mass spectrum for both proteins without lipids. We did not observe any signal intensity for the stripped monomer of the proteins, indicating both complexes were stable. Because the structures of the proteins are nearly identical, we expect the ionization efficiencies are also nearly identical between them. Any minor differences in signal intensity are thus likely due to minor errors in concentration measurements.

Next, we added lipids from the detergent-solubilized stocks. For POCL (**Figure S2–3**) and TOCL (**Figure S4–5**), we used a 1:1:50 molar ratio to achieve up to 6–7 bound lipids in the native mass spectrum and avoid overlapping of charge states observed at higher amounts of lipids. For POPG (**Figure S6–7**) and POPE (**Figure S8–9**), a molar ratio of 1:1:100 was necessary to observe enough lipids bound, indicating their weaker overall binding affinity.

Native MS of the samples was performed using a Q-Exactive HF UHMR Orbitrap mass spectrometer (Thermo Fisher Scientific, Bremen) with a variable temperature source^23^ in positive ion mode. Key settings included a mass range of 4,000−15,000 *m/z*, spray voltage of 1.2 kV, source fragmentation of 0–50 V, 75–85 V collision voltage, and a 15,000 resolution setting. Using a temperature ramp program, samples were equilibrated for 2 minutes before the acquisition of mass spectra for 1 minute from 15 to 35 °C at 5 °C intervals. We collected native MS data as single measurements from three replicate samples that were prepared separately for each mutant with each lipid.

MS data analysis was carried out using UniDec^24^ and custom Python scripts, as previously described.^6^ The deconvolution of raw mass spectra was performed using UniDec followed by 2D Grid Extraction to extract all the peak areas using the parameters described in **Table S1**. From the extracted peak areas for WT and mutant proteins with or without the lipids bound, we calculated the ratio of dissociation constant (*K*):

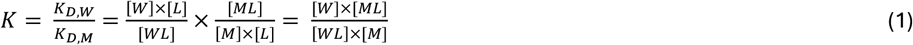

where, W, M, and L are referring to the concentration of WT protein, mutant protein, and lipid, respectively. WL and ML refer to lipid bound states of the WT and mutant protein. The equations are written as concentration for simpliciy, but native MS signal intensity is used as a proxy for concentration when calculations are made. Then, difference in free energy (*ΔΔG*) is calculated:

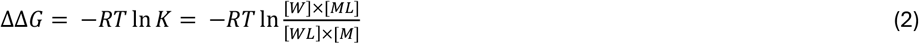

where, *R* is the ideal gas constant, *T* is the temperature in Kelvin, and *K* is the equilibrium constant from Eq. 1.

After calculating the *ΔΔG* values, we used a Python script to map the values into the Beta factor column of a PDB file of AqpZ protein. Using visual molecular dynamics (VMD)^25^ to visualize the data, we changed the graphics coloring style in the blue/white/red (BWR) method setting the midpoint at 0.5 and offset at 0. Blue indicates a favorable mutation, and red indicates an unfavorable mutation. White indicates no statistically significant effect of mutation on free energy differences. We used the values of *ΔΔG* and 95% confidence interval to calculate statistical significance. If the confidence interval crosses value 0, we consider the *ΔΔG* values to be 0 to map the residues based on free energy difference.

### Coarse-grain MD Simulations

Using the crystal structure of the wild-type AqpZ protein (PDB ID: 1RC2),^8^ we embedded the protein in a 100 by 100 Å bilayer using the CHARMM-GUI membrane builder with the Martini 22p force field.^26,27^ The membrane composition was defined to roughly model the *E. coli* lipid environment with 75% POPE, 20% POPG, and 5% CDL2 (CL) lipids in both the outer and inner leaflets.^11,28^ Note, CDL2 has similar but slightly longer tail to POPE and POPG due to one additional bead per fatty acid and includes one unsaturation per tail. We added 0.15 M sodium chloride ions and set the pH to 7.0.^26^ Four separate replicates of the AqpZ membrane bilayer were prepared.

We performed 30 µs of coarse-grain MD simulations on these membrane systems using GROMACS (version 2022.5).^29,30^ First, the system underwent an initial short energy minimization step with steepest descent algorithm, followed by a series of equilibration steps. During equilibration, various restraints on water, ions, and lipid molecules were gradually released to relax the uncorrelated initial system.^20^ An initial equilibration simulation was performed for each at 303.15 K and 1 bar with a 20 fs timestep for 3 µs. Final production simulations were run using same parameters for 30 µs. A similar procedure was followed to generate 4 replicate runs.

We calculated the occupancy and residence time of all three lipids—CL, POPE, and POPG— at each amino acid residue in the production simulations using the PyLipID package.^31^ Because AqpZ is a homo-tetramer, the occupancy and residence time values at each residue were averaged across all 4 chains and across the four replicate measurements, and the standard deviations were calculated and propagated (see SI **Tables S3** and **S4** for selected residues and Supporting Data for all residues). Based on these standard deviations, we calculated the confidence intervals at the 95% level and zeroed the values which are statistically insignificant. Using a Python script, we set the average occupancy or average residence time values as the Beta value in the AqpZ PBD file and visualized the simulations with VMD.^25^ In VMD, we changed the graphics coloring style in the blue/white/red (BWR) method setting the midpoint at 0 and offset at 0.

#### All-atom MD Simulations

To investigate the bound poses of CL on AqpZ, atomistic bilayers were prepared using the CHARMM-GUI membrane builder with the CHARMM36 force field. As with the coarse-grain simulations, we inserted AqpZ into a 100 by 100 Å *E. coli* model bilayer consisting of 75% POPE, 20% POPG, and 5% POCL. All-atom MD simulations were carried out on NAMD 3.0alpha software.^32,33^ Following a series of equilibration steps, final productions were run for 100 ns for each membrane bilayer system across three replicate runs. PyLipID analysis was performed on simulations to determine bound poses to observe lipid binding sites. We picked an example bound pose for each lipid that represents one of the sites with the highest residence time. For POPE, there were many bound poses with similar residence times, so two representatives were selected.

## Results and Discussion

Previously, we developed a single and double mutant analysis approach with native MS to assess the thermodynamic effects of perturbing specific amino acids on CL binding to AqpZ.^6^ Here, our goal was to extend this method to screen a variety of different residues around the protein-lipid interface of AqpZ to test how these mutations affected binding of different lipids. We mutated, expressed, and purified a broad range of AqpZ mutants, each containing an alanine substitution at a selected amino acid residue on the AqpZ surface (**Figure 1A–B**). These mutants included bulky hydrophobic residues that could interact with lipid tails, such as tryptophan, tyrosine, and phenylalanine. We also mutated cationic residues that could interact with lipid headgroups, such as lysine and arginine. Finally, we conducted single mutant native MS analysis^6^ on the thirteen different mutants (**Figure 1A-B**) with four different lipid types (POCL, TOCL, POPG, and POPE) (**Figure 1C–F**) to explore the site selectivity and lipid specificity of AqpZ.

### Mutant Scanning Native MS with POCL

First, we conducted single mutant experiments with POCL (**Figure 1C**) to measure the ΔΔG values for lipid binding to the mutant relative to the wild type (**Figure 2A**). A positive ΔΔG indicates decreased lipid binding to the mutant protein (unfavorable mutation). A negative value indicates enhanced lipid binding to the mutant (favorable mutation). Statistically insignificant values considering the 95% confidence interval crossing zero, indicate that the mutation had no impact on lipid binding, which could mean that either the mutated residue does not contribute to lipid binding or that the alanine substitution is not substantial enough to alter lipid binding.

**Figure 2.**
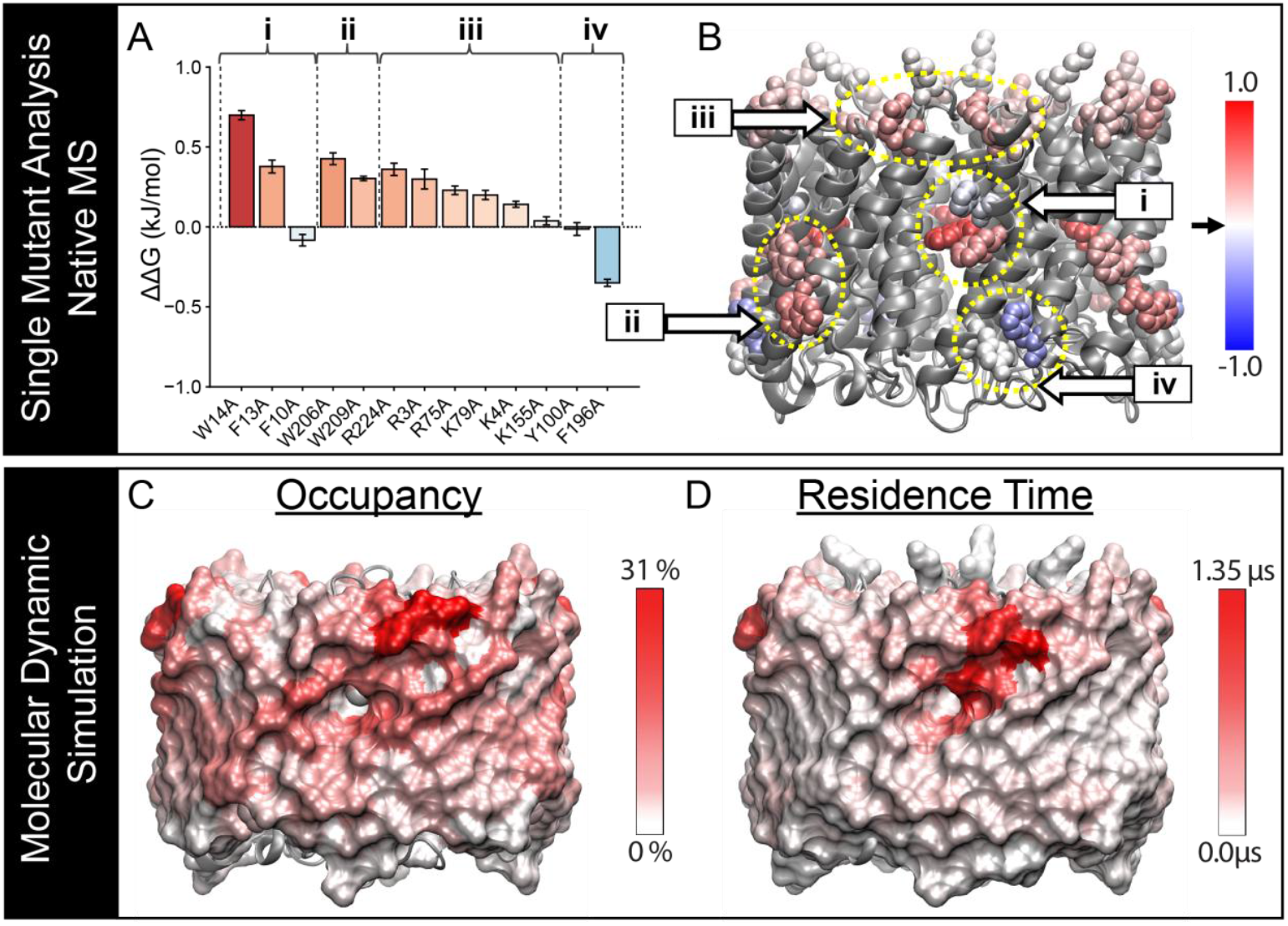
Illustrations of AqpZ-POCL interactions. (A) Difference in Gibbs free energy change (ΔΔG) for the first bound POCL at 25 °C for each mutant studied, clustered according to the identified regions. *Red* bars represent positive ΔΔG values, with the shade indicating its magnitude. *Blue* bars represent negative ΔΔG values, whereas *white* bars indicate statistically insignificant interactions. (B) AqpZ-lipid affinity maps developed based on single mutant analysis data using native MS, colored from *red* (positive ΔΔG) to *blue* (negative ΔΔG), with *white* representing no change. AqpZ-lipid affinity maps developed using MD data from (C) CL occupancy, colored from *white* (indicating no interactions) to *red* (indicating strong interactions), and (D) CL residence time, colored from *white* (indicating no interactions) to *red* (indicating stable interactions). The cytoplasmic face is shown on the top.

To highlight the hotspots of lipid binding, we mapped these ΔΔG values onto the protein structure and colored residues from red (positive ΔΔG) to blue (negative ΔΔG), with white representing no significant change (**Figure 2B**). Note, our data generated ΔΔG values for multiple steps of lipid binding, but we focused on the first lipid bound because it had the greatest magnitude of effects. The second and third bound lipid gave similar profiles but with lower magnitudes (**Figure S10**), likely due to a decrease in number of available binding sites with increased lipid binding. We did not explore cooperativity in binding here, but we expect that the binding is independent between symmetric binding sites, as was previously observed by Schmidt *et al*.^11^ We then grouped these residues into four potential binding sites based on their location, type of interaction, and ΔΔG values (regions **i**–**iv**).

We will first consider region **i**, comprised of three hydrophobic residues (W14A, F10A, and F13A) forming a binding pocket at the monomeric interface. Embedded inside the pocket (**Figure 1A-B**), W14A had the most substantial positive energy change at 0.69±0.03 kJ/mol (**Figure 2A–B** and **Table S2**), as previously observed.^6^ Prior native MS titrations showed a comparable but higher ΔΔG of around 2.6 kJ/mol for CL to the W14A mutant relative to the C20S mutant (calculated from the ratio of K_D_s), potentially due to different lipid tails used or different experimental conditions.^11^ The adjacent F13A mutation was less pronounced but still had a statistically significant ΔΔG value of 0.35±0.04 kJ/mol. F10A had little impact on lipid binding. Given the hydrophobic nature of these residues and the significant effects of the W14A and F13A mutations, it is likely that the acyl chains of the POCL molecule insert into the pocket and interact with this region. A previous cryo-EM study^15^ also indicates that this hydrophobic pocket is a potential binding site where parts of the acyl chains of CL nestle, albeit in a different orientation than we propose below.

Region **ii** consists of more exposed hydrophobic residues, W206 and W209, which may form hydrophobic interactions with lipids. Notably, both W206A and W209A mutations resulted in positive ΔΔG values of 0.43±0.04 kJ/mol and 0.30±0.01 kJ/mol, respectively (**Figure 2A–B** and **Table S2**). Thus, POCL tails may interact with these exposed and bulky hydrophobic residues.

Region **iii** consists of a cluster of cationic residues—K4, K79, K155, R3, R75, and R224—that could interact with the anionic CL headgroup (**Figure 1A and S11**). Here, we observed that three arginine mutants at the monomer-monomer interface, R224A, R3A, and R75A, had significant positive ΔΔG values, losing 0.23–0.36 kJ/mol of POCL binding energy when mutated to alanine (**Figure 2A–B** and **Table S2**). Two lysine mutants, K79A and K4A, had slightly less positive ΔΔG values (0.14–0.20 kJ/mol), and K155 was insignificant, which is consistent with its location toward the center of the protein and away from the protein-lipid interface. Thus, a subset of these residues likely interacts with the CL headgroup.

Finally, we examined F196 and Y100 in region **iv**, located on the periplasmic side of AqpZ. The same cryo-EM study suggested that the headgroup of CL is located at the hydrophilic/hydrophobic interface of the periplasmic-facing leaflet, potentially interacting with the F196 and F100 residues. However, unlike the other ΔΔG values, the F196A mutant showed a significant negative ΔΔG value of -0.35±0.02 kJ/mol (**Figure 2A–B**, and **Table S2**). Thus, mutation of the phenylalanine to an alanine enhanced binding of POCL. In contrast, Y100A had an insignificant ΔΔG value, indicating that the substitution of tyrosine with alanine at this position had no measurable effect on POCL binding. These observations suggest changes to region **iv** do not significantly disrupt POCL binding and can enhance it. Overall, these native MS data reveal how mutations to different types of residues in different regions affect POCL binding to AqpZ.

### Insights on AqpZ-CL Interactions Using MD Simulations

To gain insights into these native MS data, we performed a series of coarse-grain MD simulations of the wild-type protein placed in a membrane bilayer with 75% POPE, 20% POPG, and 5% CL. Analyzing the simulations with PyLipID,^31^ we evaluated the occupancy (**Figure 2C**) and residence time (**Figure 2D**) of CL and other lipids. Occupancy calculates the percentage of frames in which each residue contacts a target lipid.^31,34^ Residence time calculates the average length of time each residue contacts a target lipid, indicating stable interactions as bound lipids experience restricted diffusion.^31^

The occupancy data (**Figure 2C** and **Table S3**) of AqpZ-CL interactions revealed that the residues from three key regions, **i**–**iii**, demonstrate significant contacts with CL, with region **iii** (cytoplasmic cationic residues) showing the highest occupancy. Compared to other regions, region **iv** (F196 and Y100) showed a low percentage of frames that had contacts with CL (**Figure 2C** and **Table S3**).

Visualization of CL residence time on AqpZ (**Figure 2D** and **Table S4**) highlights the binding pockets of CL on AqpZ. Here, regions **i** and **iii** have stable interactions with CL, suggesting these regions could be potential CL binding sites on the inner leaflet of the membrane. This observation agrees with the expected orientation of cardiolipin on the inner leaflet.^13,35^

Although the occupancy results are comparatively higher in region **ii**, the residence time visualizations suggest that they do not stably bind with CL in this region.

Looking closely at region **i**, the F10 and F13 residues have significant occupancy (15–20%, **Table S3**), but W14 has low occupancy (1.37±1.04%, **Table S3**), despite presenting the most significant contribution in the native MS data (**Figure 2A**). Similarly, the residence time for F10 and F13 were higher (0.8–1.3 µs), but W14 had a lower residence time of 0.13±0.16 µs (**Table S4**). Our interpretation is that F10 and F13 directly interact with CL. In contrast, W14 does not directly interact with CL but instead may play an important role in maintaining the binding pocket, providing a wedge to hold open the site for F10 and F13 to interact. Interestingly, F10 is seen strongly interacting in the MD data, but the F10A mutation does not significantly affect POCL binding, indicating that the alanine interacts similarly with this set of lipid tails as phenylalanine in this site.

To gain molecular insights into our proposed CL binding orientation of AqpZ based on native MS data, we constructed an all-atom model of the wild-type protein placed in a similar membrane bilayer and simulated for 100 ns using NAMD. Next, we analyzed the simulations using the PyLipID python package to extract bound poses of POCL.^31^ In line with the native MS data, we observed a binding conformation where the lipid tails interact predominantly with the hydrophobic pocket in region **i** at the monomeric interface and the headgroup interacts with positively charged residues on the cytoplasmic interface in region **iii** (**Figure 3**). This binding pose had the highest residence times, highest surface area of contacts, and lowest RMSD, indicating it is likely one of the preferred orientations. It also agrees with prior molecular dynamics simulations on this system.^11,13^

**Figure 3.**
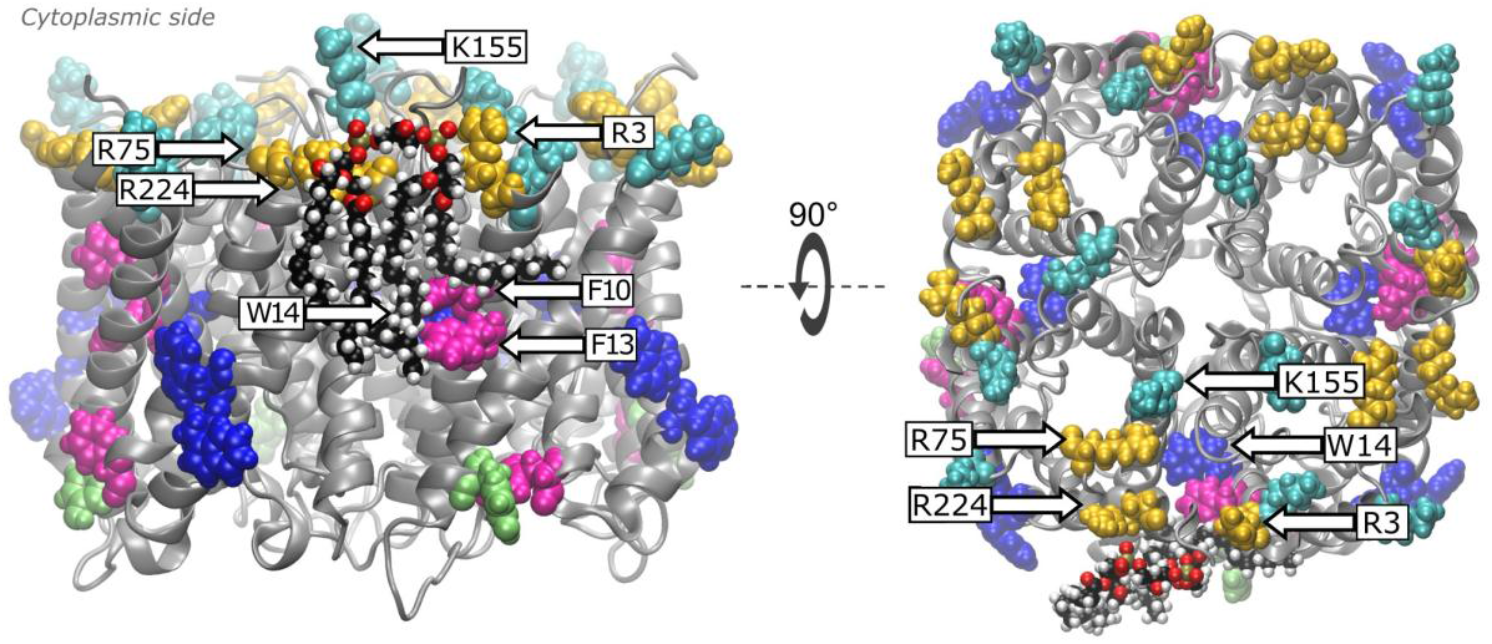
Side view (*left*) and top view (*right*) of a representative POCL bound pose determined using PyLipID for the binding site 1 from an all-atom MD simulation of AqpZ wild-type protein in a bilayer mimicking *E. coli* lipids. The protein is colored in *gray*, and the selected residues are colored in their respective color as described in Figure 1. The CL lipid is shown in a ball-and-stick representation colored by atom name. The top view structure (right) is oriented with cytoplasmic face in front.

Additionally, another binding pose was found where the POCL tail interacts with the hydrophobic region **ii** (W206 and W209) and faces the periplasmic surface **(Figure S12)**. However, this orientation was not consistently observed across replicates and generally had lower residence times, lower surface areas, and higher RMSD values than the top binding pose. Thus, this pose suggests that region **ii** is not the highest affinity POCL binding site but may be a secondary lipid binding site. A previous cryo-EM structure^15^ also proposed a periplasmic orientation but had a different lipid position than seen here. Together, these data suggest that a lower affinity site could be present on the periplasmic side.

### Selectivity of Different Lipids for AqpZ Binding Sites

After studying AqpZ interactions with POCL at different binding sites, we next explored the selectivity of these mutations towards different lipids using native MS. First, we investigated TOCL (**Figure 1D**), which has the same headgroup but different acyl chains. Then, we tested POPG (**Figure 1E**) and POPE (**Figure 1F**), which have the same tails but different headgroups.

#### Effect of Lipid Tails

TOCL is similar to POCL but with all four acyl chains unsaturated (18:1) (**Figure 1D**), which makes TOCL likely more fluid in bilayer environments and less tight when packing.^36^ We performed similar single mutant native MS analyses for each mutant with TOCL. The ΔΔG plot shows many similarities and some differences compared to POCL (**Figure 4**).

**Figure 4.**
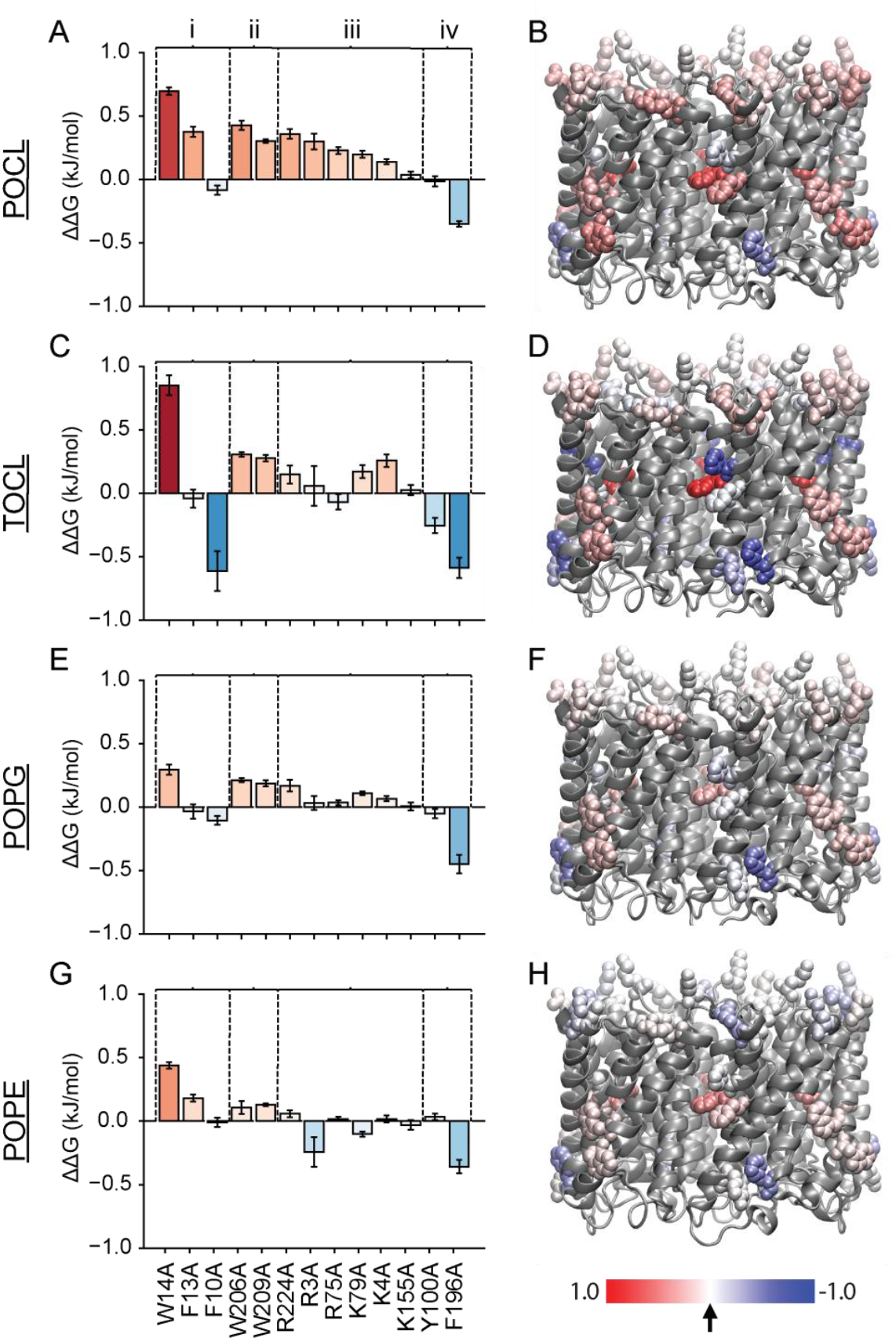
Single mutant analysis data using native MS for (A, B) POCL, (C, D) TOCL, (E, F) POPG, and (G, H) POPE. *Left* panel indicates ΔΔG changes for the first bound lipid at 25 °C for each mutant studied, clustered according to the identified regions. *Red* bars represent positive ΔΔG values. *Blue* bars represent negative ΔΔG values, and *white* bars indicate statistically insignificant interactions. *Right* panel indicates AqpZ-lipid affinity maps, colored from *red* (positive ΔΔG) to *blue* (negative ΔΔG), with *white* representing no change. Panels A and B are reproduced from Figure 2 for comparison. Structures are oriented with the cytoplasmic face on top.

Considering region **i** (F10, F13, W14), the W14A mutation that had the most significant impact with POCL also had the highest contribution with TOCL (**Figure 4C–D** and **Table S2**). However, the contribution of F13A became insignificant, and the small ΔΔG observed for F10A with POCL became significantly lower negative ΔΔG (p-value=0.003, **Figure 4C** and **Table S2**). Here, the F10A mutation now shows a significant negative ΔΔG, indicating that the less bulky alanine is more favorable than phenylalanine for binding TOCL in this site. The difference in F10A response between POCL and TOCL reveals how lipid tails can interact differently with the protein surface in the presence of different mutants. These effects indicate that the acyl chain significantly affects interactions to hydrophobic residues in region **i**, consistent with the structure in **Figure 3**.

Although the ΔΔG values are slightly different, region **ii** (W206 and W209) is not dramatically affected by lipid tail differences between TOCL and POCL (**Figure 4C** and **Table S2**). Similarly, in region **iii**, mutants R75A (p-value=0.0008), R224A (p-value=0.0066), and K4A (p-value=0.001) exhibit statistically significant differences in TOCL binding compared to POCL, but R3A, K79A, and statistically significant differences in TOCL binding compared to POCL, but R3A, K79A, and K155A show no significant differences (**Figure 4C** and **Table S2**). As above, K155 serves as a negative control that likely does not interact with lipids. Overall, lipid tail differences between TOCL and POCL do not dramatically affect interactions in region **iii**, which are instead likely driven by headgroups.

In region **iv**, the ΔΔG values for Y100A and F196A became more negative from POCL to TOCL. Both Y100A (p-value=0.005) and F196A (p-value=0.003) mutants are more favorable to TOCL binding compared with POCL (**Figure 4C** and **Table S2**). Together, these data indicate that CL binding in some sites is unaffected by lipid tails, particularly with cationic residues in region **iii**. However, the differences in the acyl chain do affect how the lipid interacts with some hydrophobic binding regions, especially **i** and **iv**.

#### Effect of Lipid Headgroups

Next, we investigated whether these AqpZ sites are specific for CL headgroups compared to other lipid types. To study this lipid specificity, we used phospholipids with similar acyl chains (16:0_18:1) but with different headgroups, including anionic PG (**Figure 1E**) and zwitterionic PE (**Figure 1F**). By comparing the ΔΔG values and maps generated for each lipid (**Figure 4**), we observed a significant reduction (p-values<0.05) in the absolute value of ΔΔG for almost all the mutants with POPG and POPE compared to POCL (**Figure S13**). This overall reduced ΔΔG indicates that the mutations cause the greatest disturbance for CL binding compared to PG and PE, which suggests that these residues showed higher specificity towards CL over other lipids.

First, we compared POCL with POPG to evaluate the effects of mutations on two anionic lipids with similar structures. Here, the ΔΔG values were comparable but reduced in magnitude in regions **i, ii**, and **iv**. In region **iii** (the cytoplasmic cationic residues), the effect of all mutations was reduced compared to POCL, but a few retained positive ΔΔG values. Thus, POPG can be affected by mutation of some of the positively charged amino acid residues on the cytoplasmic interface, but mutations to these sites affect CL binding more dramatically.

To test the effects of headgroup charge, we compared anionic POCL against zwitterionic POPE. Here, the ΔΔG trends were comparable between POCL and POPE in regions **i, ii**, and **iv**, albeit with lower overall magnitudes of ΔΔG with POPE. However, in region **iii**, most residues showed insignificant ΔΔG values for POPE, indicating that interactions with the positive amino acid residues are not substantial. This observation further confirms that region **iii** has a significant influence on lipid headgroup interactions through salt bridge interactions with anionic lipids like PG and CL.

Overall, mutations to specific AqpZ residues affect CL more dramatically compared to both POPG and POPE (**Figure S13**). Mutants in regions **i** and **iii** are more affected by lipid headgroup than other potential binding regions. Furthermore, among other residues, the contributions from W14 and F196 for each lipid binding remain notable, possibly indicating a global role these residues play in the structural arrangement of the binding sites/pockets.

### Insights into Lipid Specificity of AqpZ Binding Sites Using MD Simulations

To evaluate lipid selectivity in the MD simulations, we conducted PyLipID analysis for POPE and POPG lipids using the same simulations described above. Because CL hasFresF lower abundance in the membrane (5%), the occupancy data were consistently lower compared to the other two lipids (**Figure 5**). Thus, we will focus our attention on the residence time data.

**Figure 5.**
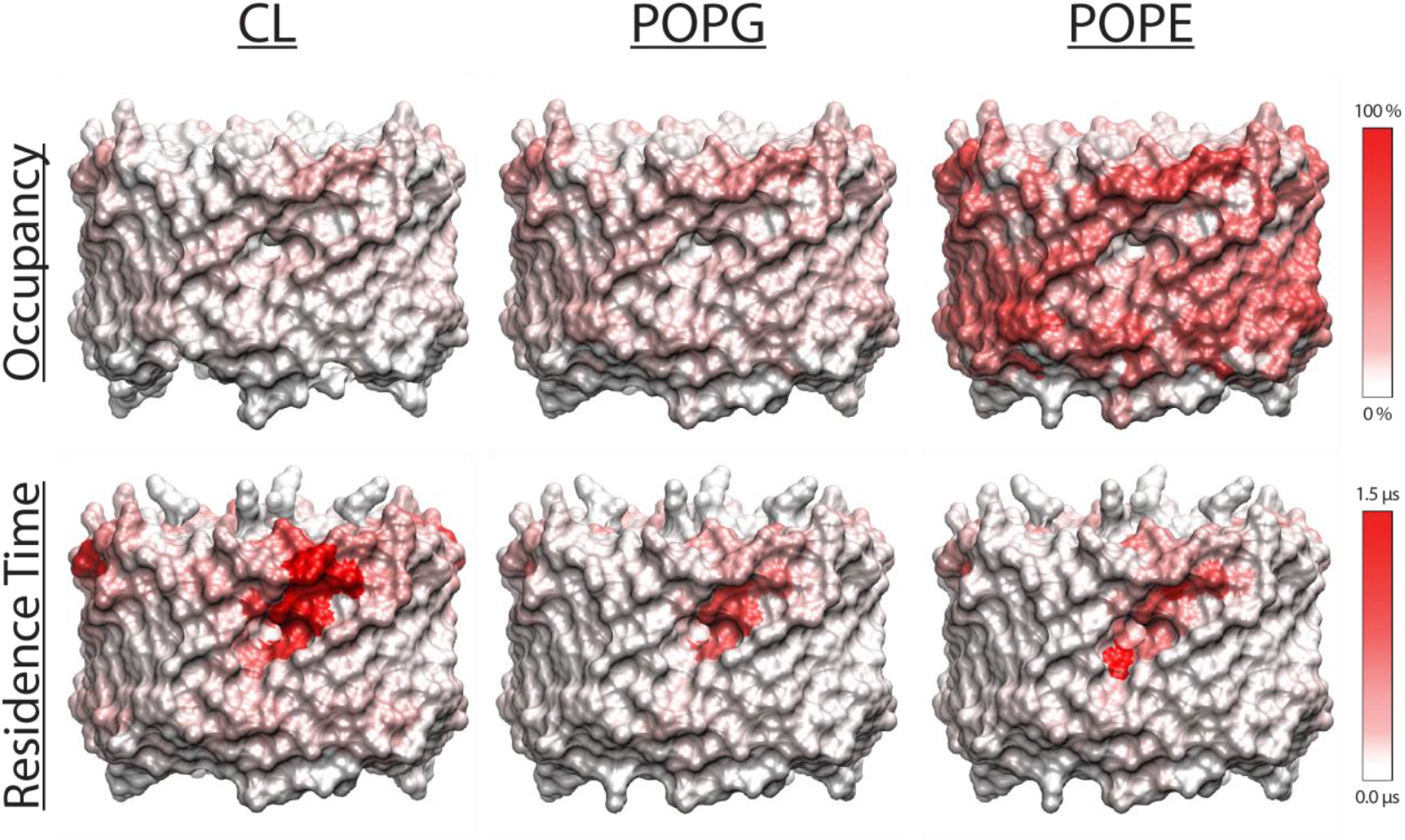
Coarse-grain MD illustrations of AqpZ-lipid interactions. The average occupancy (*top*) and average residence time (*bottom*) at selected residues on different wild-type AqpZ by CL (*left*), POPG lipids (*middle*), and POPE lipids (*right*). The maps are colored from *white* (indicating no interactions) to *red* (indicating strong/stable interactions). Structures are oriented with the cytoplasmic face on top.

The residence time data clearly demonstrates that between these three lipids studied, CL had significantly higher (p-value<0.05) residence times at many of the selected residues compared to POPG and POPE. Thus, CL forms the most kinetically stable interactions with AqpZ (**Figure 5**), which is consistent with our native MS data and confirms that AqpZ shows selectivity towards CL in these sites.

Considering the location of lipid binding, each lipid type tends to interact more stably in regions **i** and **iii** compared to regions **ii** and **iv (Figure 5** and **Table S4)**. This observation suggests that the site formed by regions **i** and **iii** may serve as a lipid binding site. In contrast, regions **ii** and **iv** show comparatively small residence times for all three lipids (less than 0.1 µs), indicating relatively weak lipid binding ability. Bound pose analysis of POPG and POPE in all-atom MD simulations showed a number of different binding poses for each lipids. Some are at the monomer interface around region **i** and **iii** (**Figures S14** and **S15A, B**). Others are located at other sites around the protein surface (**Figure S15C, D**). For both POPG and POPE, a greater diversity of binding poses was observed than with POCL.

Combining both native MS and MD simulation observations, we found that AqpZ forms more stable interactions with CL relative to PG and PE, which is consistent with the CL binding in previous research using lipid exchange-MS,^9^ native MS titrations,^11^ and prior molecular dynamics studies.^11,13^ Looking closely at tail interactions and headgroup interactions, it appears that interactions with hydrophobic residues at the hydrophobic pocket in region **i** and the electrostatic interactions with the cytoplasmic cationic residues in region **iii** are the most important interactions for lipid stability.

## Conclusions

Here, we profiled the lipid binding sites of AqpZ and explored its lipid specificity, creating a lipid binding map through an integrated approach combining native MS and MD simulations. Our data revealed that AqpZ binds to CL with the greatest specificity, and it is less specific for PG and PE. When CL binds to AqpZ in its tightest location, the CL acyl chains likely interact with the hydrophobic pocket at the monomeric interface with the headgroup orienting towards cationic residues at the cytoplasmic surface of AqpZ. This site was selective for CL and the primary site with stable lipid binding.

Our screening of single mutants revealed that the W14 residue plays a crucial role in lipid binding. Interestingly, W14 does not seem to directly interact with the lipid itself, but it likely facilitates the structural arrangement of a binding pocket that interacts with CL. Thus, residues may facilitate the formation of lipid-binding sites without direct interactions. These observations provide new insights into membrane protein-lipid interactions that can be captured through experimental techniques but can be challenging to capture in simulations alone. Together, combining the results from both native MS and MD enabled us to achieve a broader perspective on AqpZ lipid binding sites and their specificity.

## AUTHOR INFORMATION

### Conflict of Interest

The authors declare no competing financial interest.

### Supporting Information

The Supporting Information includes supplemental tables with data analysis results, and supplemental figures on single mutant analysis MS data (raw and deconvolved) and affinity maps of AqpZ; AqpZ structures; comparison of energetics between different lipids; and lipid bound poses. Raw mass spectrometry data for this manuscript is available at MassIVE MSV000096149 (DOI: 10.25345/C5SJ1B35F). Full MD results are provided as a Supporting Data file.

## ACKNOWLEDGMENT

The authors thank Maria Reinhardt-Szyba, Kyle Fort, and Alexander Makarov at Thermo Fisher Scientific for support on the Q-Exactive HF UHMR instrument. We used High Performance Computing (HPC) resources supported by the University of Arizona TRIF, UITS, and Research, Innovation, and Impact (RII) and maintained by the UArizona Research Technologies department. We also thank Megan Ewbank for early work on mutagenesis experiments and Levi Brown and Marius Kostelic for making a variable temperature source in house. This work was funded by the National Institutes of Health (NIH) under grant numbers R35 GM128624 and RM1 GM145416 to M.T.M.

## ACCESSION CODES

AqpZ: UniProt P60844

## For Table of Contents use only

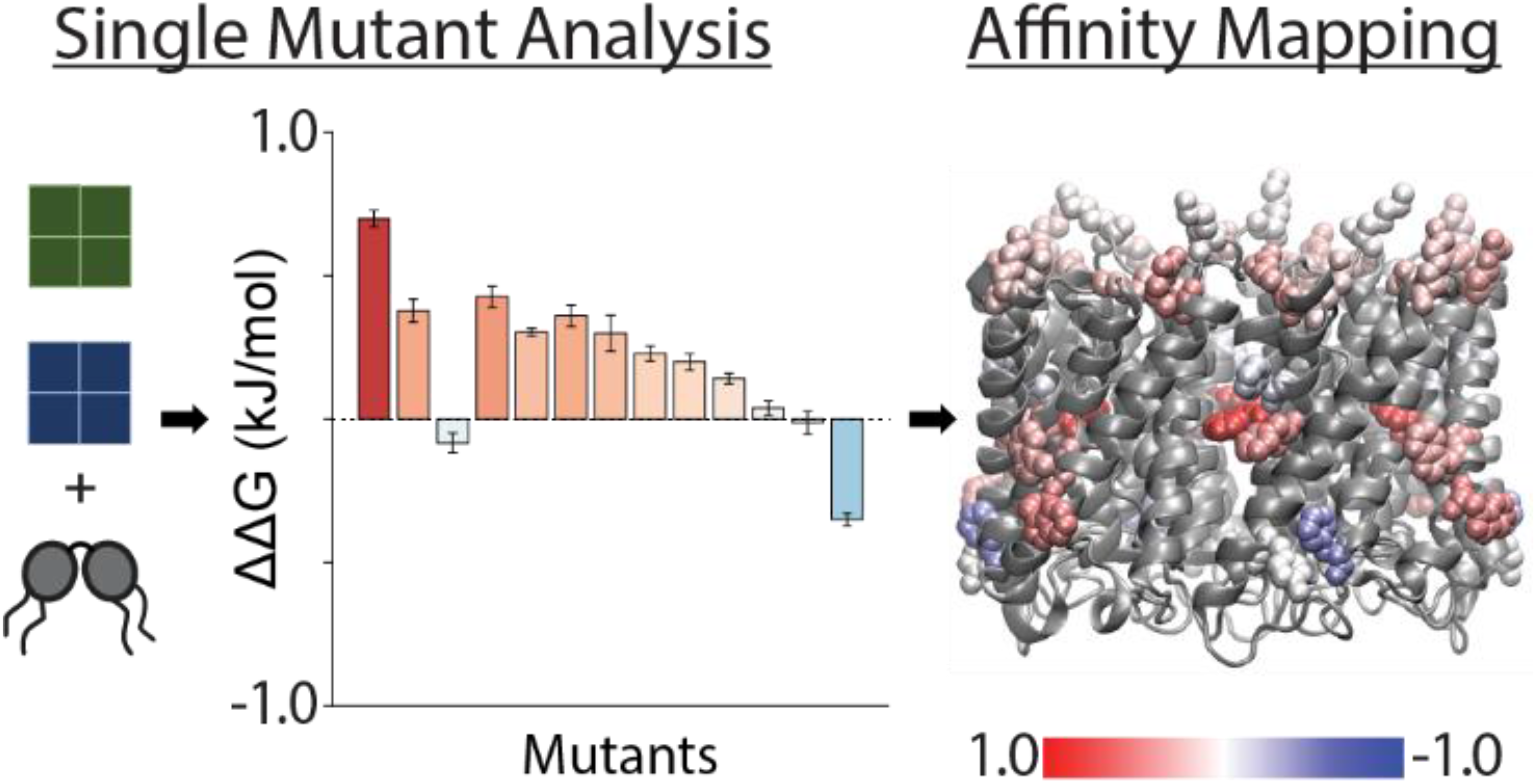

## Notes

https://doi.org/doi:10.25345/C5SJ1B35F

### Competing Interest Statement

The authors have declared no competing interest.

### Summary of Updates

Revisions include new figures, expanded discussions, and clarifications to figures and text.

