## Supporting Information for "Alanine Scanning to Define Membrane Protein-Lipid Interaction Sites Using Native Mass Spectrometry"

TABLE OF CONTENTS

Supplemental Tables....................................................................................................................2

Supplemental Figures....................................................................................................................6

### Supplemental Tables

**Table S1**: Parameters used in extracting the deconvolved mass peaks using 2D grid extraction tool in UniDec. Mass 0 corresponds to the mass of the membrane protein with the lower mass (the mutant protein), mass 1 is the mass of lipids, and mass 2 corresponds to the mass difference between the WT protein (98940 Da) and the mutant protein. Each mass value represents an input into the UniDec extraction tool.

| Membrane Protein A | Membrane Protein B | Mass 0 (Da) | Mass 1 (Da) | | | | Mass 2 (Da) |
|---|---|---|---|---|---|---|---|
| | | | POCL | TOCL | POPG | POPE | |
| WT | R3A, R75A, R224A | 98600 | 1405 | 1458 | 747 | 718 | 340 |
| WT | K4A, K79A, K115A | 98700 | 1405 | 1458 | 747 | 718 | 240 |
| WT | F10A, F13A, F196A | 98570 | 1405 | 1458 | 747 | 718 | 370 |
| WT | W14A, W206A, W209A | 98445 | 1405 | 1458 | 747 | 718 | 495 |
| WT | Y100A | 98545 | 1405 | 1458 | 747 | 718 | 355 |

**Table S2**: Single Mutant Analysis. The mean difference in the Gibbs free energy change (ΔΔG (kJ/mol)) and the 95% confidence interval (CI) for POCL, TOCL, POPG, and POPE binding between each pairwise membrane protein combination of wild-type and each mutant for the first bound lipid at 25 °C.

| **Mutant** | **POCL** | | **TOCL** | | **POPG** | | **POPE** | |
|---|---|---|---|---|---|---|---|---|
| | Mean ΔΔG (kJ/mol) | CI | Mean ΔΔG (kJ/mol) | CI | Mean ΔΔG (kJ/mol) | CI | Mean ΔΔG (kJ/mol) | CI |
| W14A | 0.70 | 0.03 | 0.85 | 0.08 | 0.30 | 0.04 | 0.44 | 0.02 |
| F13A | 0.38 | 0.04 | -0.04 | 0.07 | -0.03 | 0.06 | 0.18 | 0.03 |
| F10A | -0.08 | 0.04 | -0.61 | 0.16 | -0.10 | 0.03 | -0.01 | 0.04 |
| W206A | 0.43 | 0.04 | 0.31 | 0.02 | 0.21 | 0.02 | 0.11 | 0.05 |
| W209A | 0.30 | 0.01 | 0.28 | 0.03 | 0.19 | 0.02 | 0.13 | 0.01 |
| R224A | 0.36 | 0.04 | 0.15 | 0.07 | 0.17 | 0.05 | 0.06 | 0.03 |
| R3A | 0.30 | 0.06 | 0.22 | 0.13 | 0.03 | 0.06 | -0.24 | 0.12 |
| R75A | 0.23 | 0.03 | -0.07 | 0.06 | 0.04 | 0.02 | 0.02 | 0.02 |
| K79A | 0.20 | 0.03 | 0.17 | 0.05 | 0.11 | 0.01 | -0.10 | 0.02 |
| K4A | 0.14 | 0.02 | 0.26 | 0.05 | 0.07 | 0.02 | 0.02 | 0.03 |
| K155A | 0.04 | 0.02 | 0.03 | 0.04 | 0.01 | 0.03 | -0.03 | 0.04 |
| Y100A | -0.01 | 0.04 | -0.25 | 0.06 | -0.05 | 0.04 | 0.03 | 0.03 |
| F196A | -0.35 | 0.02 | -0.59 | 0.08 | -0.45 | 0.07 | -0.36 | 0.05 |

**Table S3**: Coarse-grain MD simulation data for lipid occupancy of selected residues. The averaged occupancy and 95% confidence interval (CI) calculated for CL, POPG, and POPE lipid at each residue on AqpZ wild-type protein. This data shows an average of 4 replicate MD simulations. Data was averaged from the 4 simulations and 4 binding sites within each simulation (n=16). The standard deviation was propagated, and the confidence interval was calculated at the 95% confidence level. Additional data for every residue is provided in the Supplemental Data.

| Residue | CL | | POPG | | POPE | |
|---|---|---|---|---|---|---|
| | Average Occupancy | CI Occupancy | Average Occupancy | CI Occupancy | Average Occupancy | CI Occupancy |
| 14TRP | 1.37 | 0.56 | 4.14 | 2.49 | 8.13 | 3.37 |
| 13PHE | 19.92 | 4.00 | 20.75 | 3.43 | 41.92 | 2.36 |
| 10PHE | 15.92 | 3.58 | 19.70 | 4.23 | 31.23 | 4.11 |
| 206TRP | 12.59 | 2.06 | 23.70 | 2.79 | 69.34 | 5.80 |
| 209TRP | 12.41 | 1.42 | 20.32 | 2.38 | 56.18 | 5.17 |
| 224ARG | 5.96 | 1.90 | 14.05 | 2.88 | 23.48 | 3.79 |
| 3ARG | 18.72 | 3.63 | 28.89 | 3.93 | 37.27 | 3.30 |
| 75ARG | 3.37 | 0.66 | 9.42 | 1.44 | 16.28 | 2.67 |
| 79LYS | 13.30 | 2.36 | 28.26 | 3.78 | 44.18 | 5.04 |
| 4LYS | 8.31 | 2.10 | 18.28 | 3.64 | 22.33 | 4.36 |
| 155LYS | 0.16 | 0.15 | 0.16 | 0.08 | 0.31 | 0.28 |
| 100TYR | 3.83 | 0.80 | 11.28 | 1.47 | 37.92 | 4.57 |
| 196PHE | 7.54 | 1.47 | 20.85 | 2.02 | 62.01 | 5.35 |

**Table S4**: Coarse-grain MD simulation data for the residence times of lipids. The averaged residence time and 95% confidence interval (CI) calculated for CL, POPG, and POPE lipid at each residue on AqpZ wild-type protein. Averages and 95% CI were calculated as in Table S3. Additional data for every residue is provided in the Supplemental Data.

| Residue | CL | | POPG | | POPE | |
|---|---|---|---|---|---|---|
| | Average Residence Time | CI Residence Time | Average Residence Time | CI Residence Time | Average Residence Time | CI Residence Time |
| 14TRP | 0.13 | 0.08 | 0.12 | 0.07 | 0.31 | 0.18 |
| 13PHE | 0.84 | 0.30 | 0.62 | 0.25 | 0.49 | 0.11 |
| 10PHE | 1.28 | 0.43 | 0.70 | 0.26 | 0.64 | 0.14 |
| 206TRP | 0.10 | 0.02 | 0.06 | 0.01 | 0.10 | 0.03 |
| 209TRP | 0.07 | 0.02 | 0.04 | 0.01 | 0.05 | 0.01 |
| 224ARG | 0.24 | 0.06 | 0.13 | 0.03 | 0.11 | 0.03 |
| 3ARG | 0.68 | 0.20 | 0.31 | 0.08 | 0.35 | 0.08 |
| 75ARG | 0.07 | 0.01 | 0.04 | 0.01 | 0.04 | 0.01 |
| 79LYS | 0.32 | 0.09 | 0.11 | 0.02 | 0.07 | 0.02 |
| 4LYS | 0.36 | 0.07 | 0.45 | 0.10 | 0.32 | 0.06 |
| 155LYS | 0.02 | 0.02 | 0.02 | 0.01 | 0.04 | 0.03 |
| 100TYR | 0.16 | 0.05 | 0.09 | 0.02 | 0.09 | 0.02 |
| 196PHE | 0.20 | 0.02 | 0.08 | 0.02 | 0.08 | 0.01 |

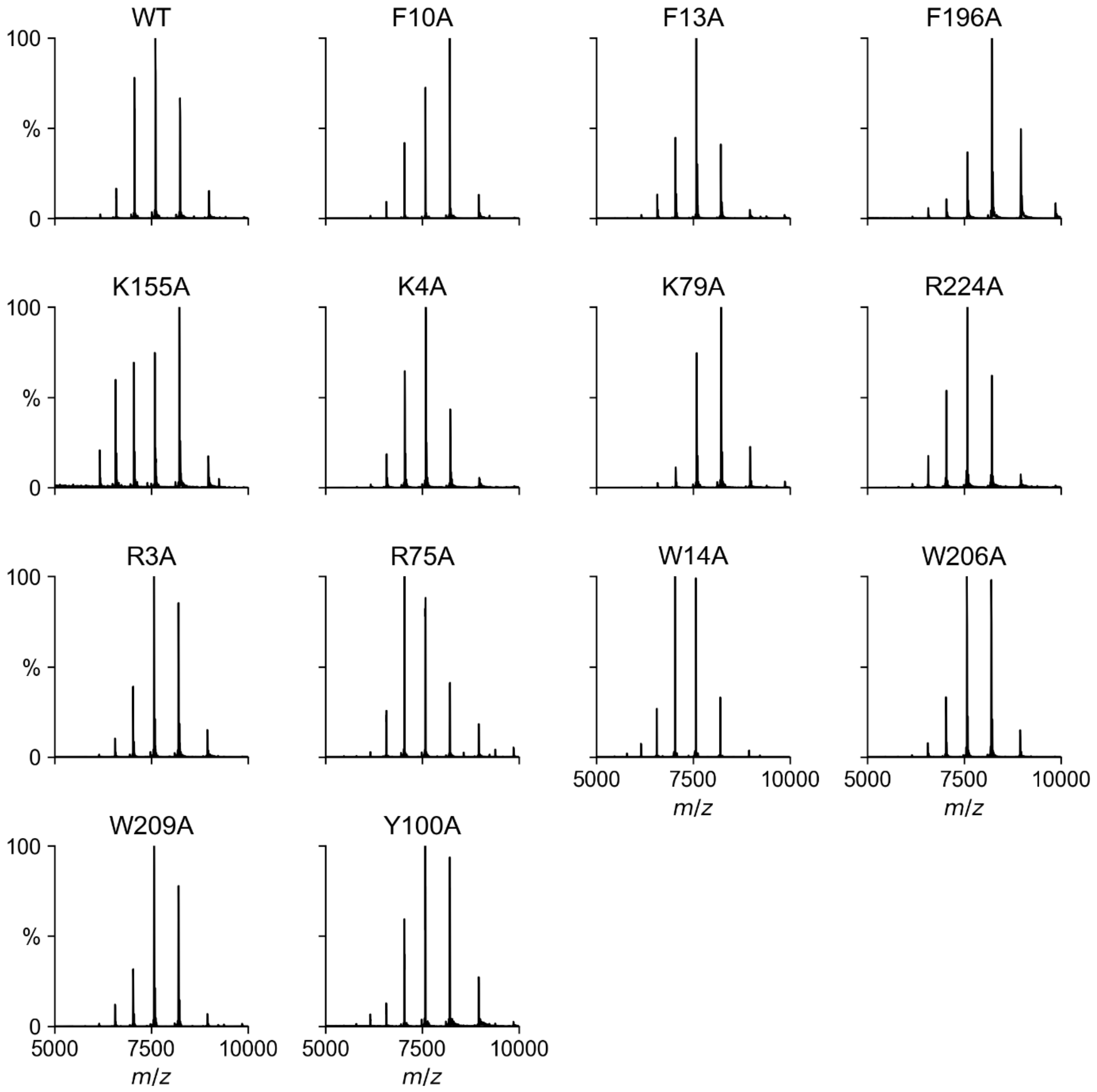


**Figure S1:** Raw mass spectra for AqpZ WT and mutant proteins. Note, these spectra are for individual proteins only, not mixtures of two species. The major peaks observed were for the tetramer charge state distribution. Minor peaks in the R75A spectrum are for a dimer of tetramers.

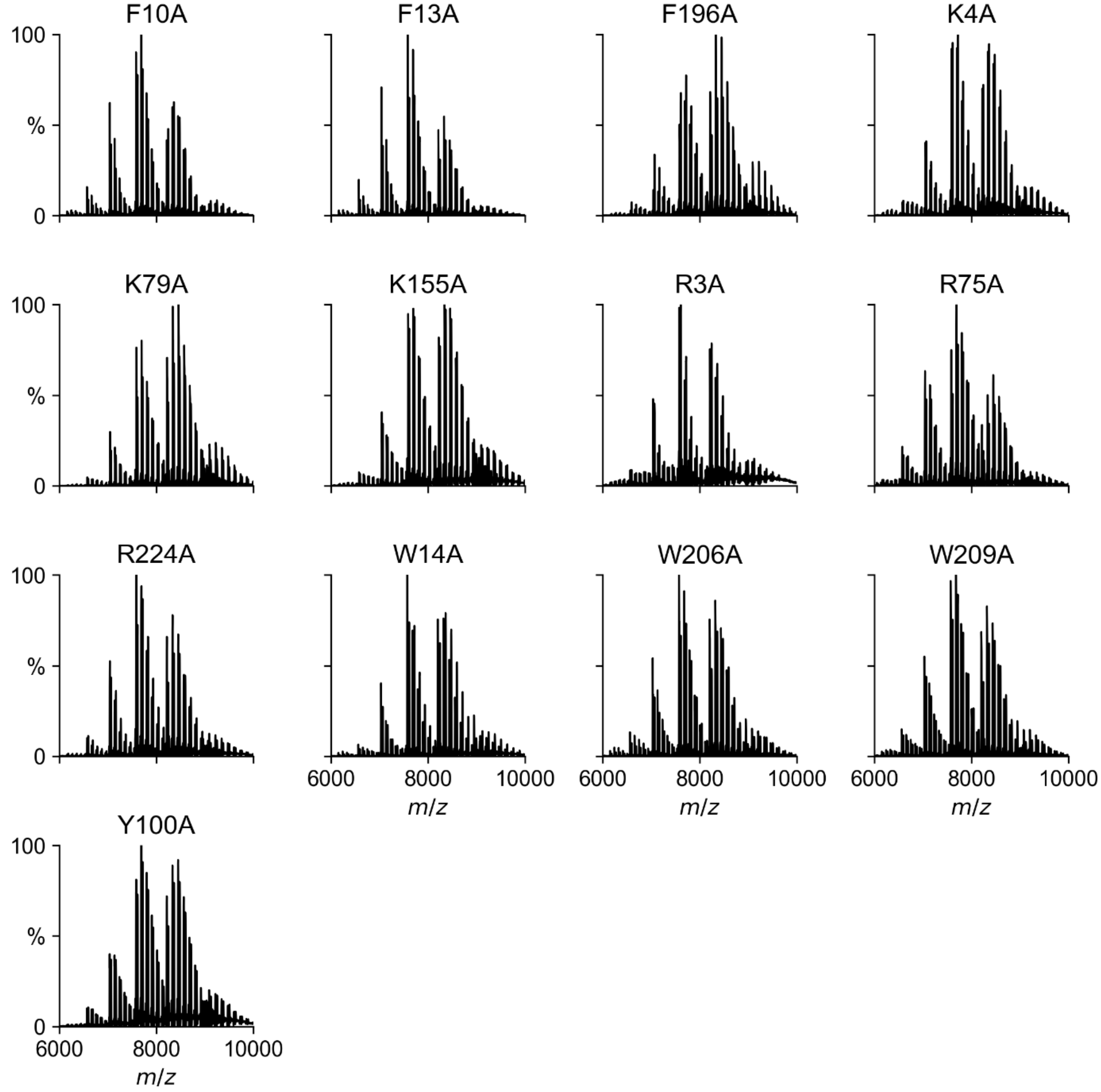


**Figure S2:** Raw mass spectra for a representative replicate of AqpZ mutants with wild-type protein bound to POCL. Spectra are averaged across all temperature steps.

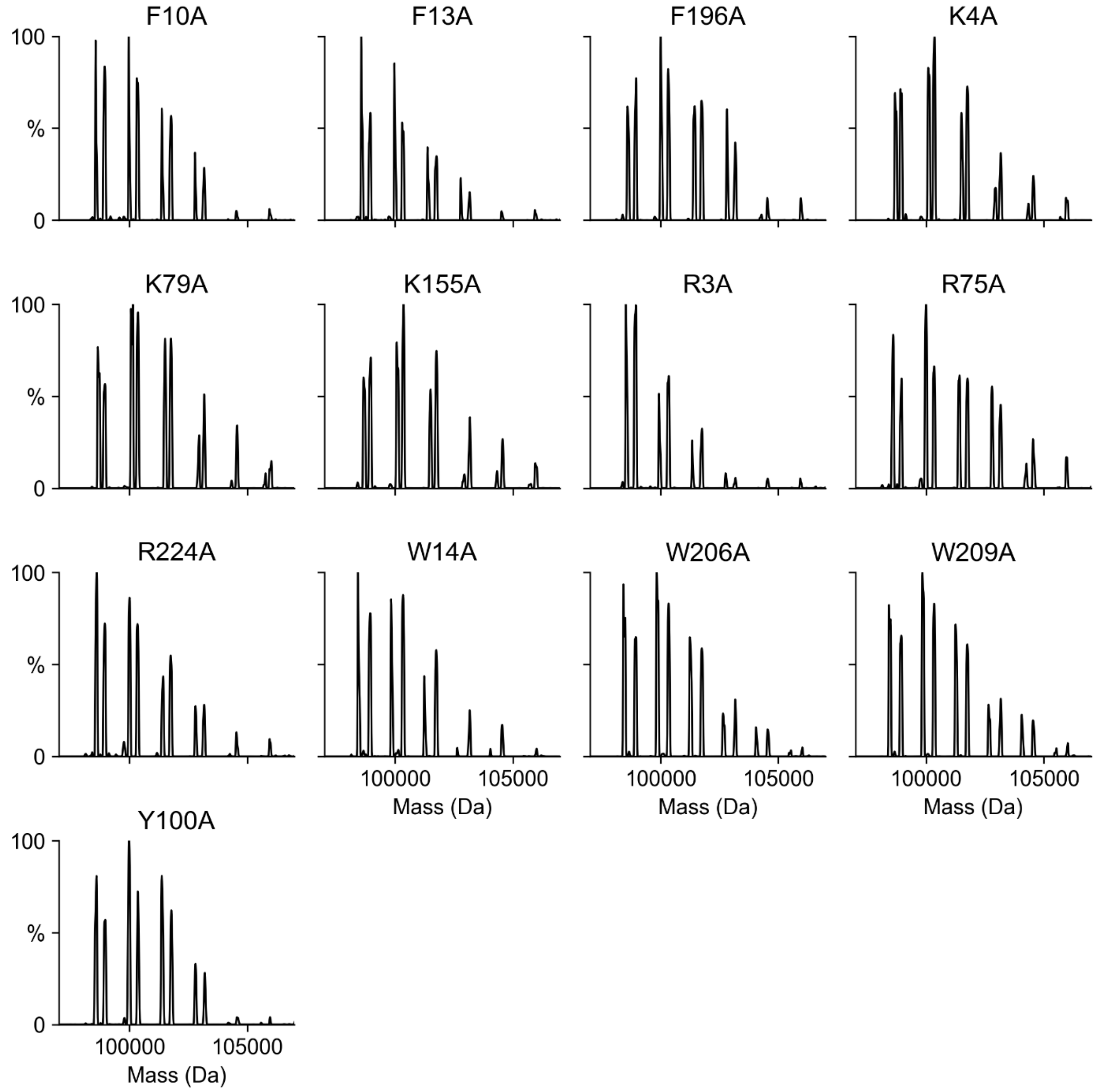


**Figure S3:** Deconvolved mass spectra for a representative replicate from Figure S2 of AqpZ mutants with wild-type protein bound to up to six lipids in complex with POCL.

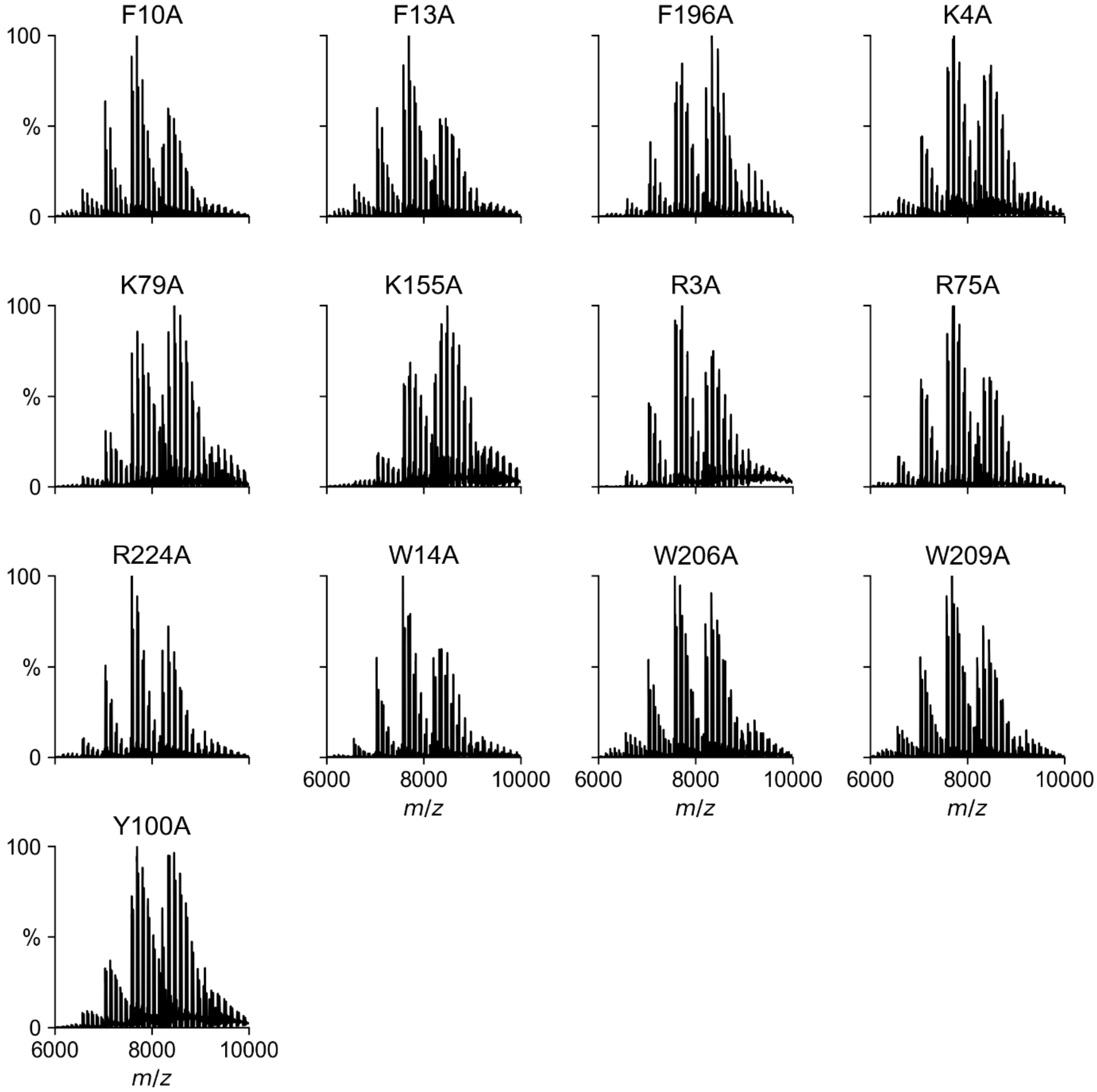


**Figure S4:** Raw mass spectra for a representative replicate of AqpZ mutants with wild-type protein bound to up to six lipids in complex with TOCL. Spectra are averaged across all temperature steps.

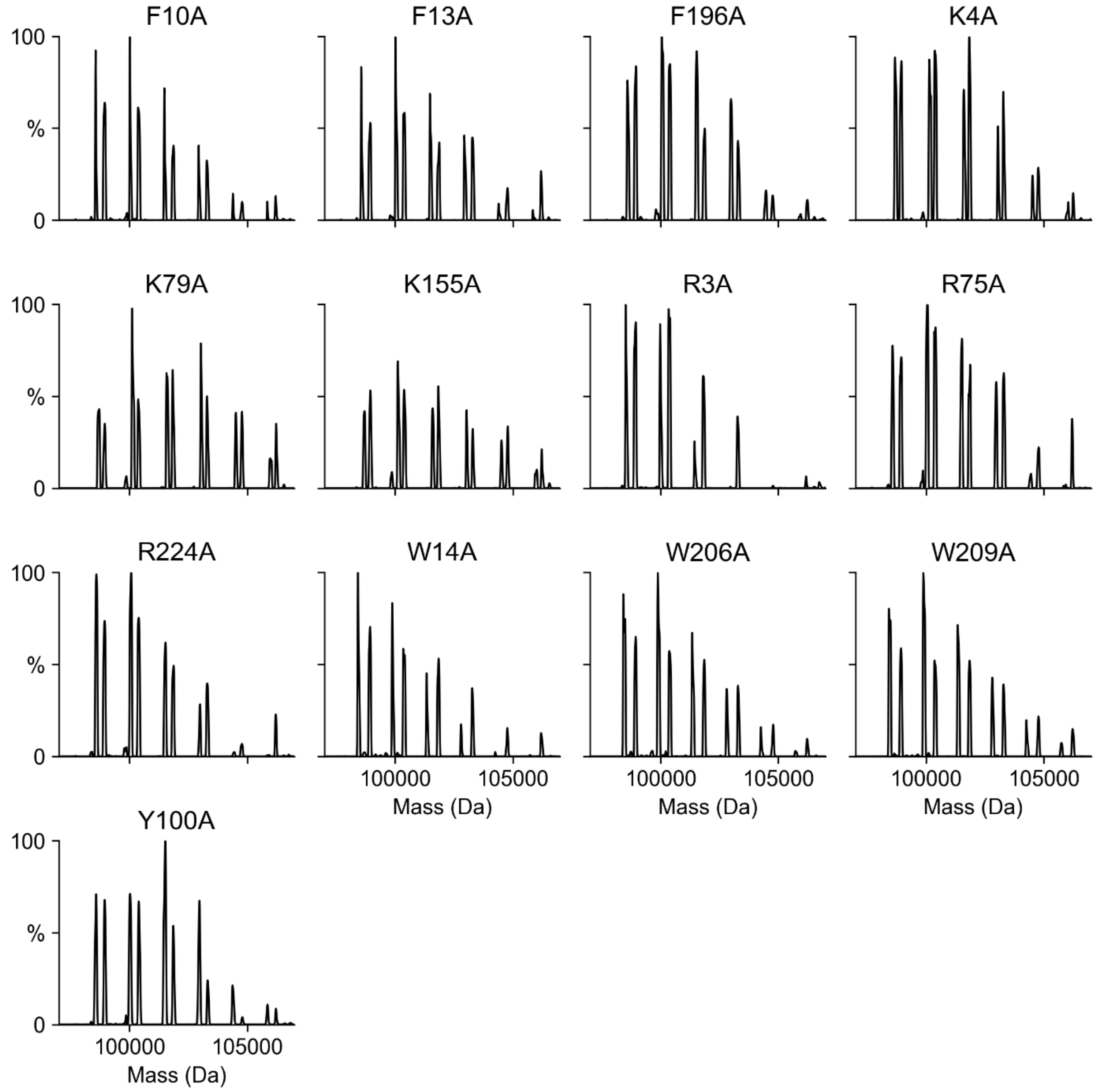


**Figure S5:** Deconvolved mass spectra for a representative replicate from Figure S4 of AqpZ mutants with wild-type protein bound to up to six lipids in complex with TOCL.

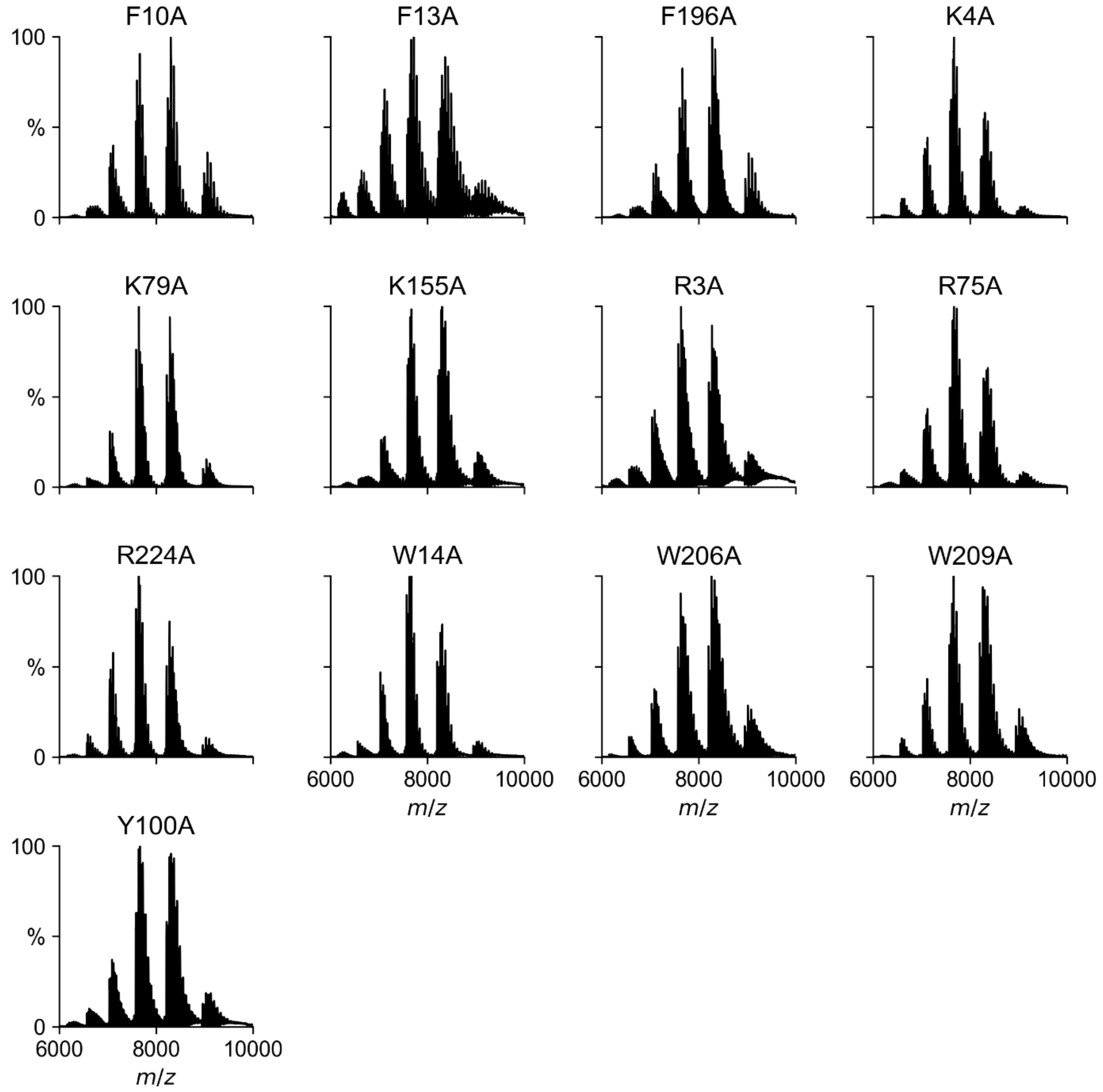


**Figure S6:** Raw mass spectra for a representative replicate of AqpZ mutants with wild-type protein bound to up to six lipids in complex with POPG. Spectra are averaged across all temperature steps.

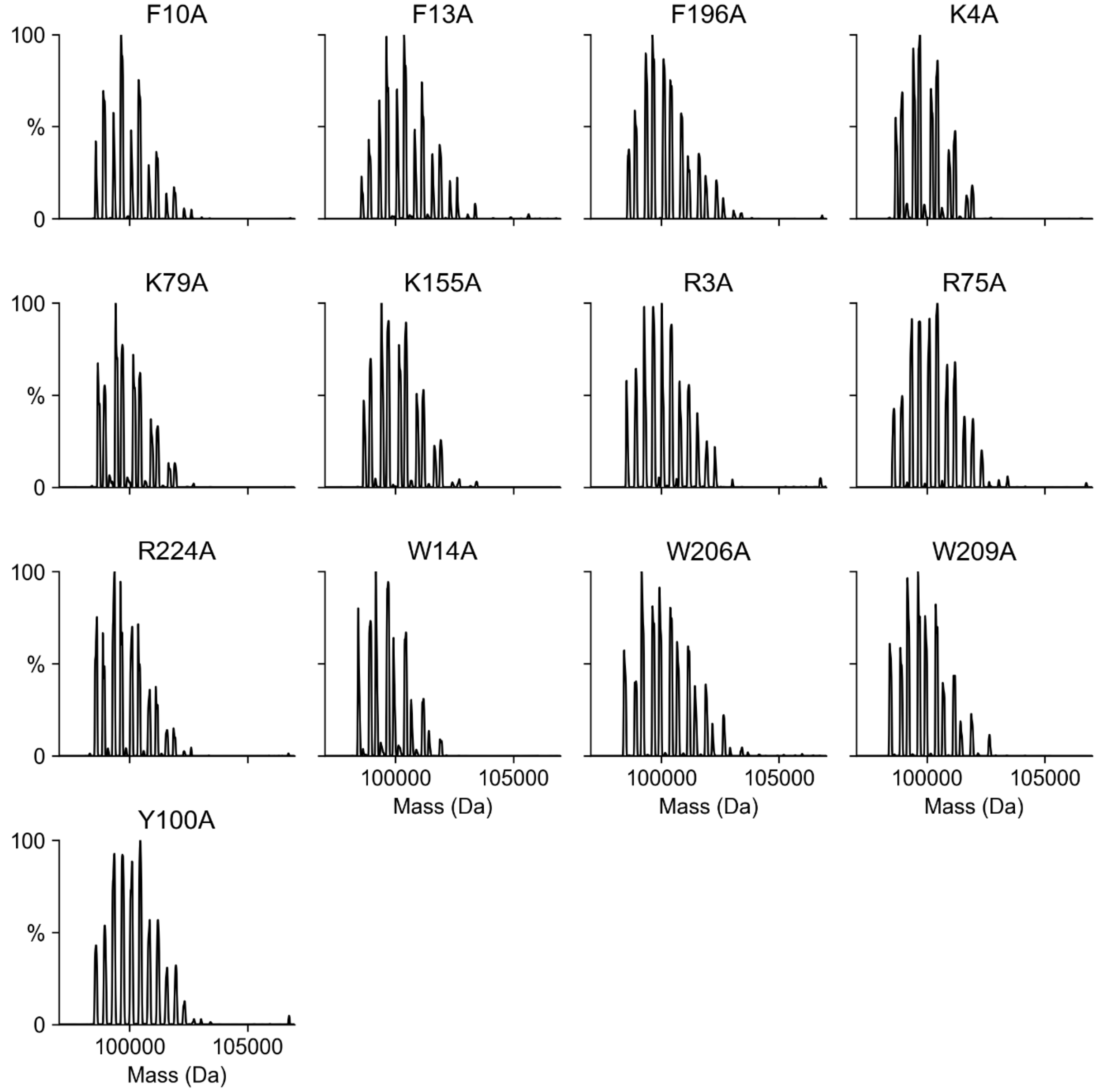


**Figure S7:** Deconvolved mass spectra for a representative replicate from Figure S6 of AqpZ mutants with wild-type protein bound to up to six lipids in complex with POPG.

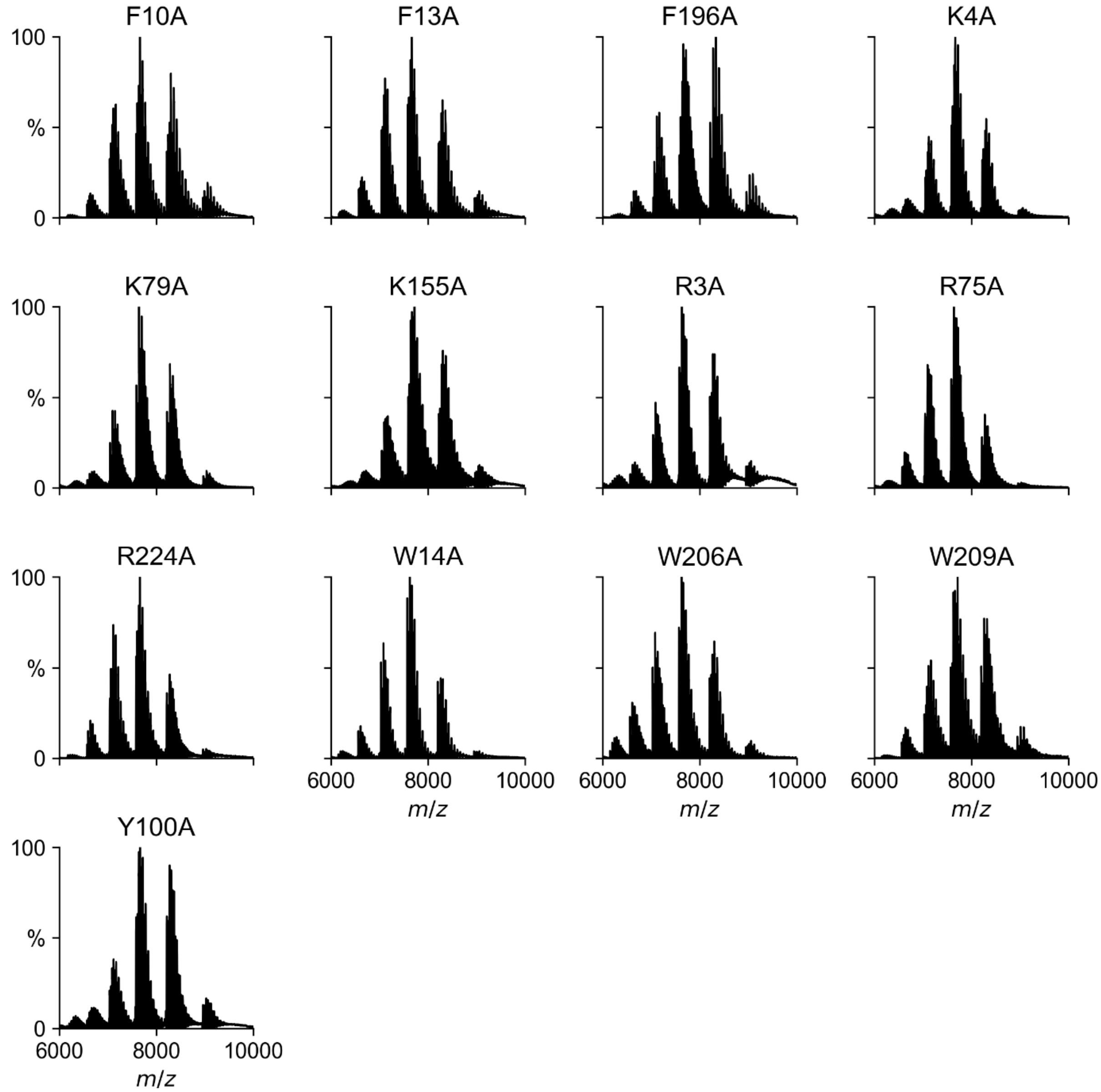


**Figure S8:** Raw mass spectra for a representative replicate of AqpZ mutants with wild-type protein bound to up to six lipids in complex with POPE. Spectra are averaged over all temperature steps.

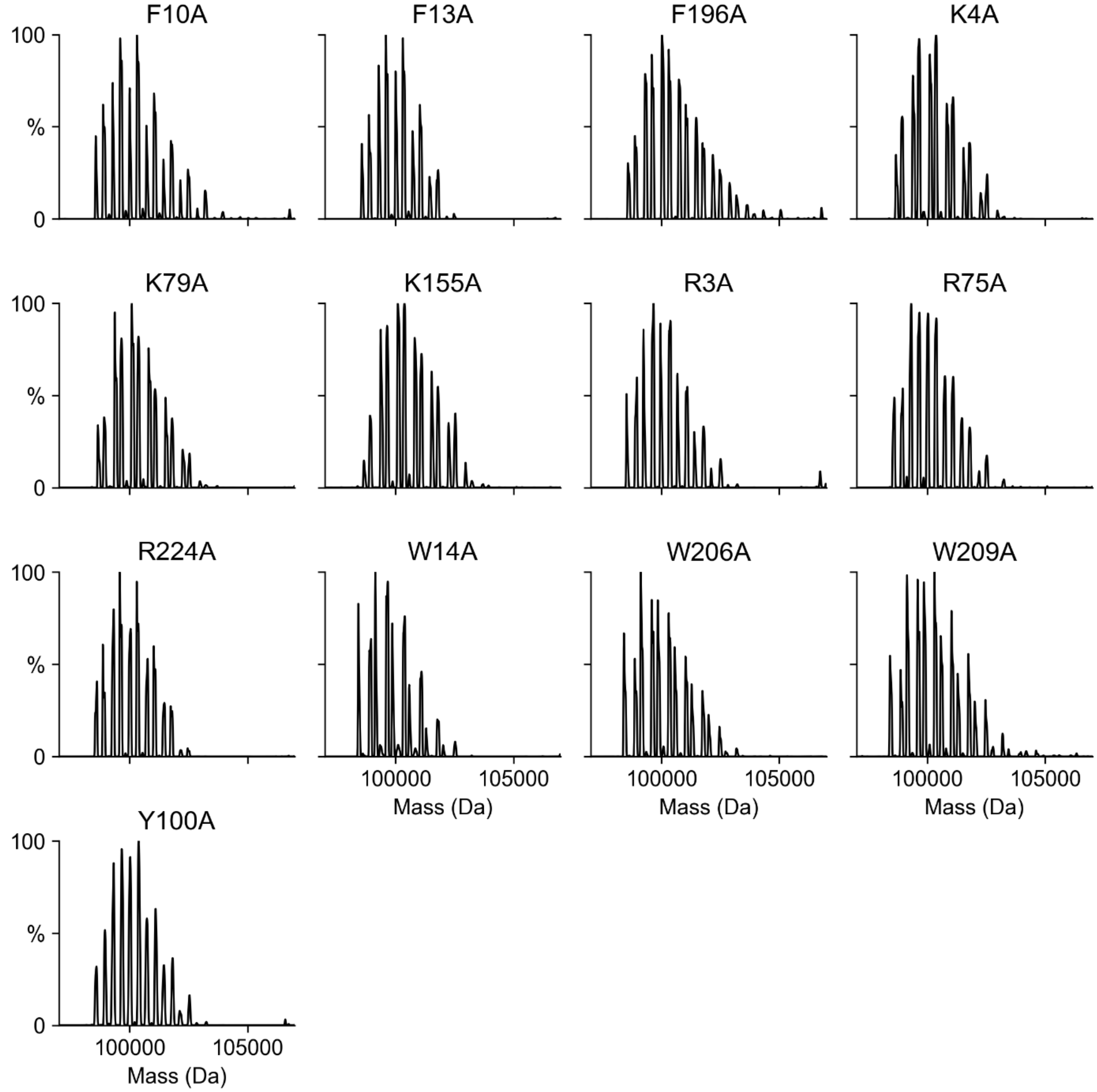


**Figure S9:** Deconvolved mass spectra for a representative replicate from Figure S8 of AqpZ mutants with wild-type protein bound to up to six lipids in complex with POPE.

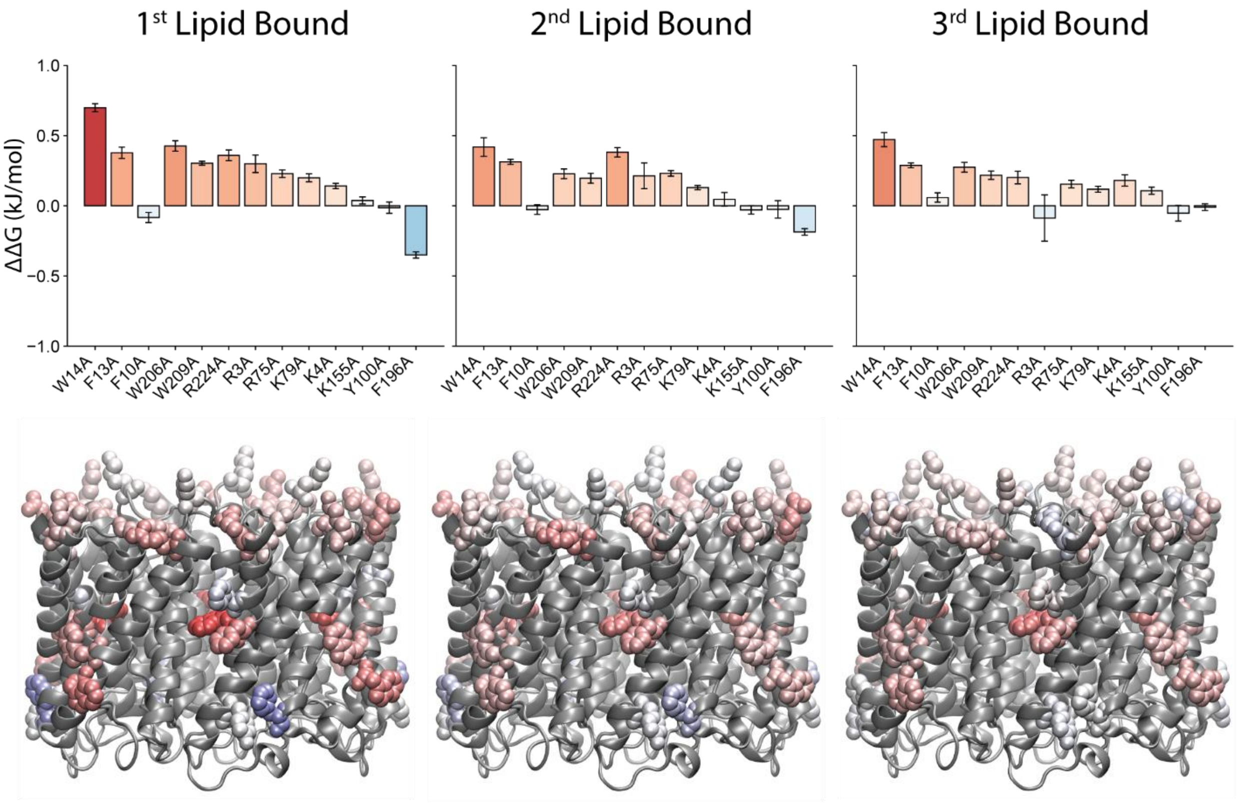


**Figure S10.** A comparison of the ΔΔG plots at 25 °C for the first, second, and third bound lipids of POCL (*top*) shows similar trends, but with decreasing magnitude for increasing lipid number. The corresponding affinity maps (*bottom*) illustrate the relative lipid binding affinity of selected residues colored from *red* (positive ΔΔG) to *blue* (negative ΔΔG), with *white* indicating no change. Structures are oriented with the cytoplasmic face on top.

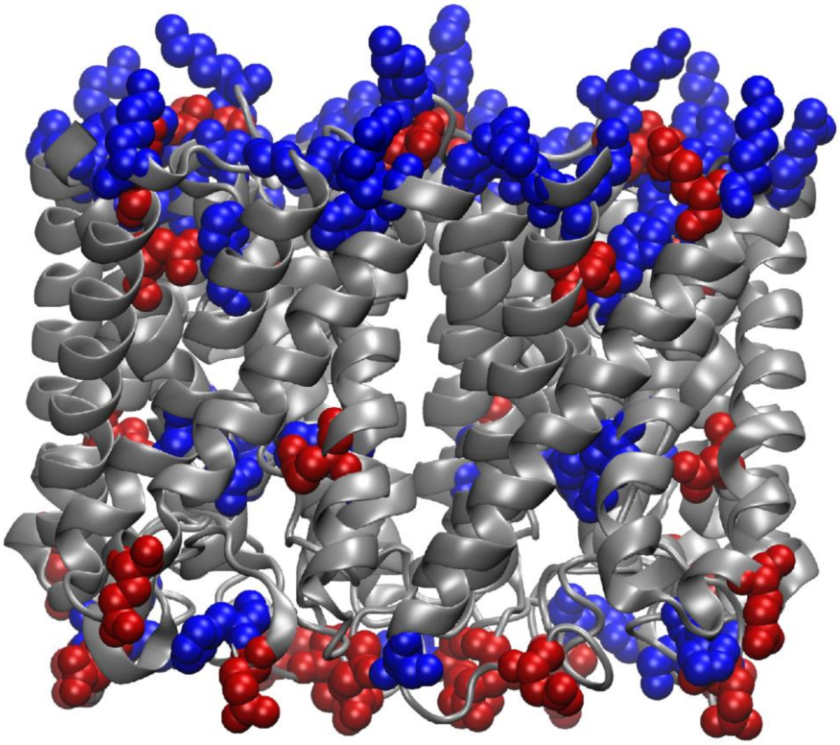

**Figure S11**. Side-view of AqpZ structure colored with acidic and basic residues shown in *red* and *blue*, respectively. The cytoplasmic side is shown as the top with the transmembrane plane in the middle and intermembrane side on bottom.

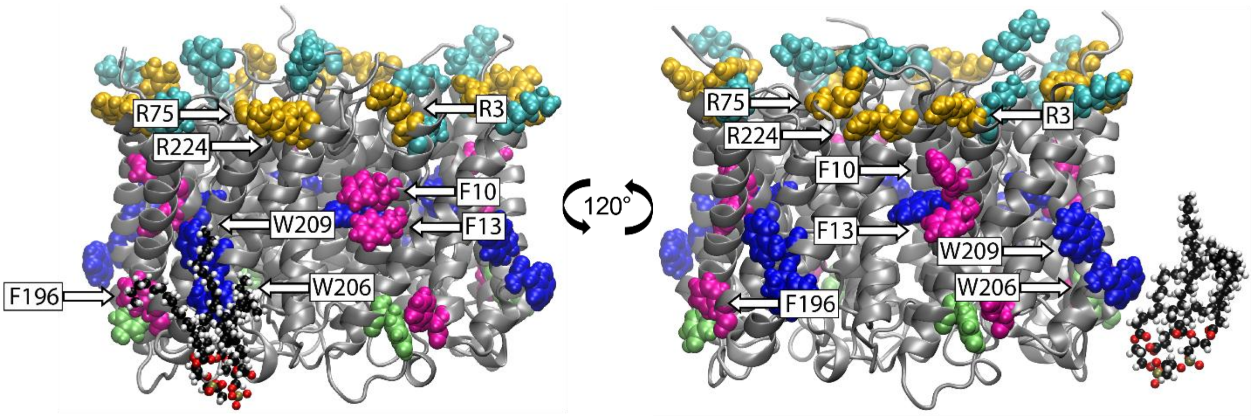


**Figure S12:** A representative structure determined using PyLipID for the second most stable binding pose of CL from an all-atom MD simulation of AqpZ wild-type protein in a bilayer mimicking *E. coli* lipids. The protein is colored *gray*, and the selected residues are colored in their respective color as described in Figure 1. The CL lipid is shown in a ball-and-stick representation colored by atom name. The primary interactions of the lipids are with W206 and W209 in region **ii**. Structures are oriented with the cytoplasmic face on top.

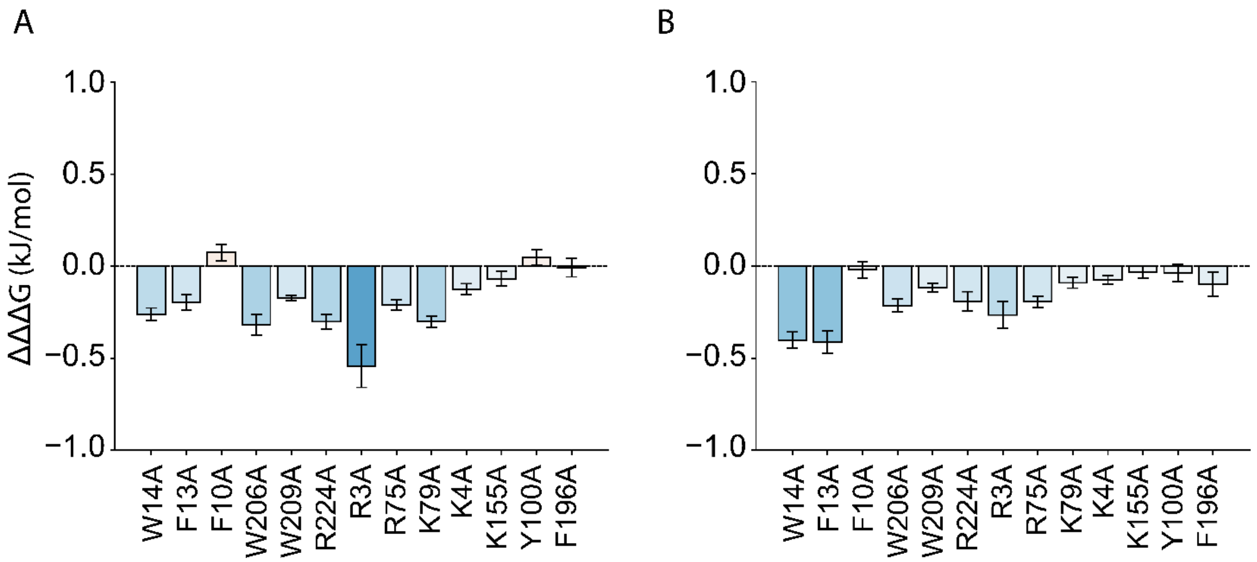


**Figure S13:** Difference in ΔΔG between (A) POCL and POPE, and (B) POCL and POPG. Negative values indicate larger ΔΔG values in POCL. Positive values indicate larger ΔΔG values in the POPE or POPG.

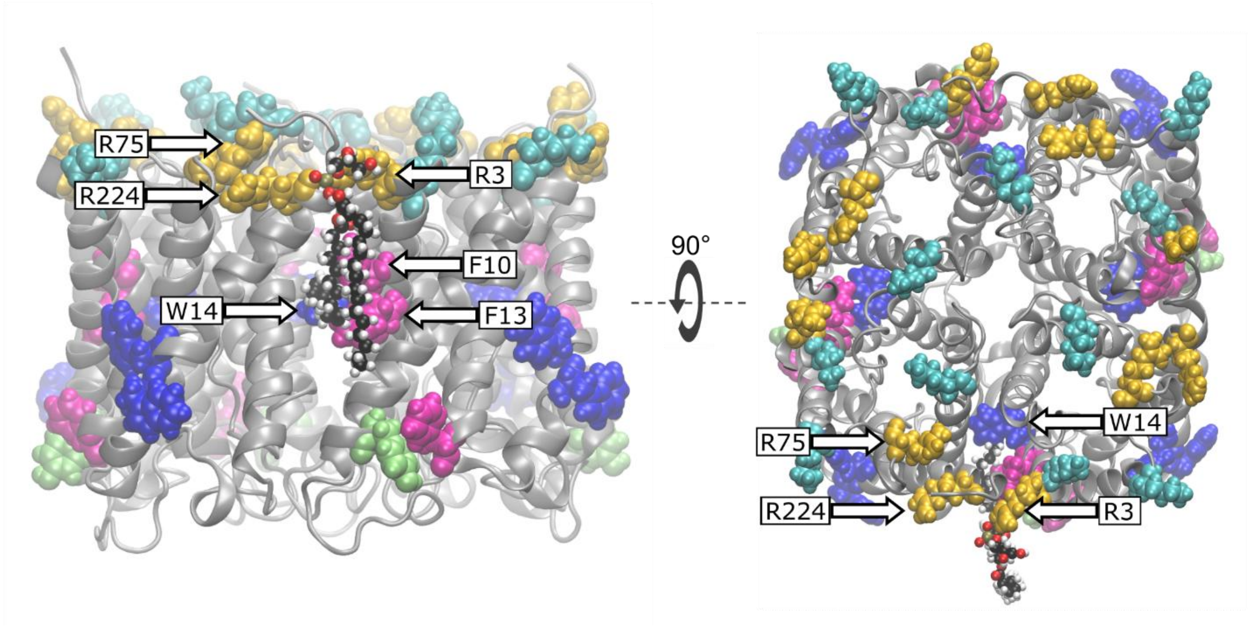


**Figure S14:** Side view (*left*) and top view (*right*) of an example POPG bound pose determined using PyLipID from an all-atom MD simulation of AqpZ wild-type protein in a bilayer mimicking *E. coli* lipids. The protein is colored in *gray*, and the selected residues are colored in their respective color as described in Figure 1. The POPG lipid is shown in a ball-and-stick representation colored by atom name. The side view structure (*left*) is oriented with the cytoplasmic face on top. The top view strcuture (*right*) is oriented with cytoplasmic face in front.

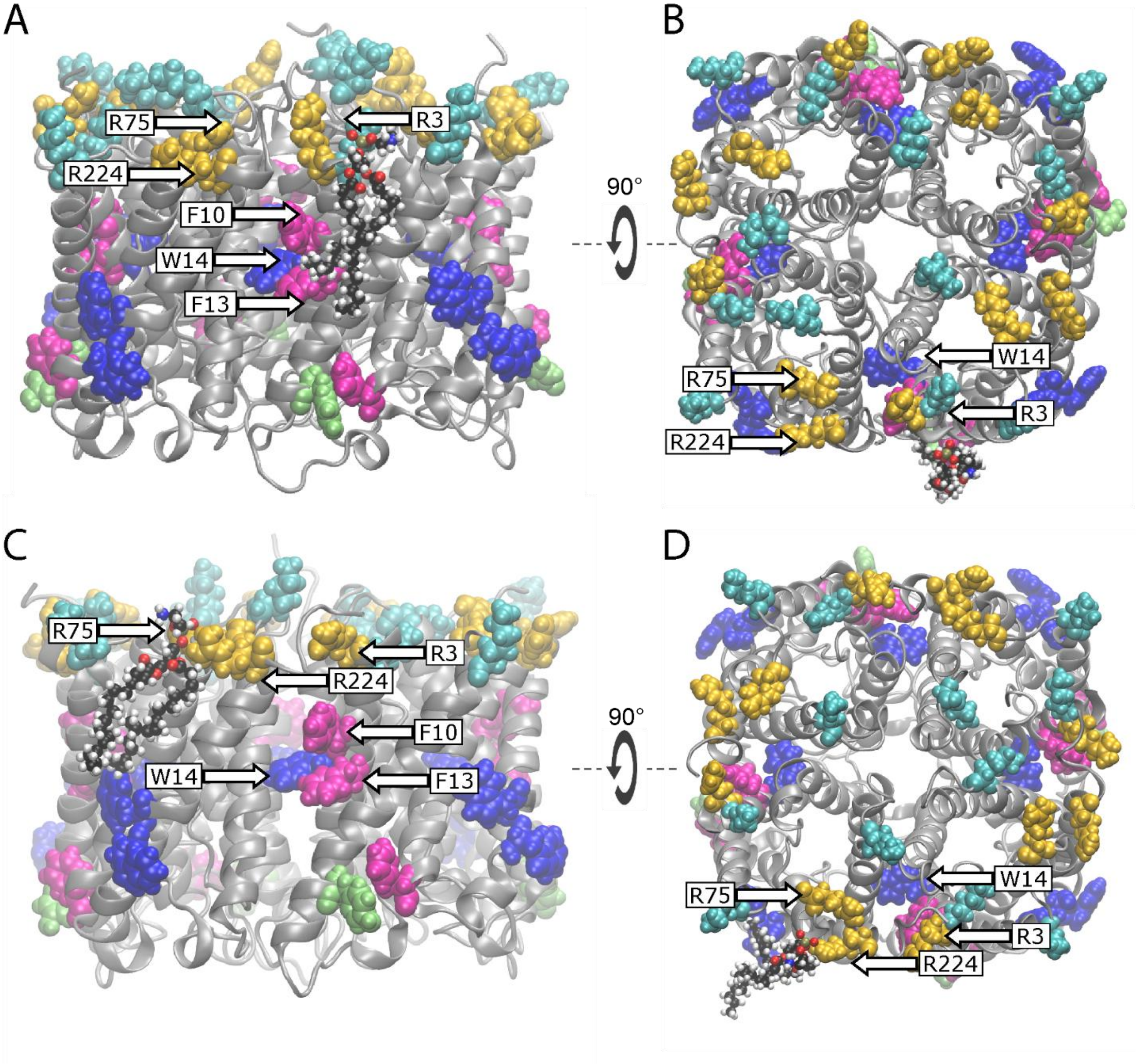


**Figure S15:** Side view (A, C) and top view (B, D) of two example POPE bound poses determined using PyLipID from an all-atom MD simulation of AqpZ wild-type protein in a bilayer mimicking *E. coli* lipids. Each bound pose is selected based on high residence time. The protein is colored in *gray*, and the selected residues are colored in their respective color as described in Figure 1. The POPE lipid is shown in a ball-and-stick representation colored by atom name. Side view structures (A, C) are oriented with the cytoplasmic face on top. The top view strcutures (B, D) are oriented with cytoplasmic face in front.
